# Temporal Analyses of Postnatal Liver Development and Maturation by Single Cell Transcriptomics

**DOI:** 10.1101/2021.07.14.451852

**Authors:** Yan Liang, Kota Kaneko, Bing Xin, Jin Lee, Xin Sun, Kun Zhang, Gen-Sheng Feng

## Abstract

Liver is the major metabolic organ, although its postnatal development and maturation are inadequately understood. We analyzed 52,834 single cell transcriptomes and identified 31 cell types or states in mouse livers at postnatal day 1, 3, 7, 21 and 56. We observed unexpectedly high levels of hepatocyte heterogeneity in the developing liver and progressive construction of the zonated metabolic functions from pericentral to periportal hepatocytes, which was orchestrated with development of sinusoid endothelial, stellate and Kupffer cells. Trajectory and gene regulatory analyses captured 36 transcription factors, including a circadian regulator Bhlhe40, in programming liver development. Remarkably, we identified a special group of macrophages enriched at day 7 with a hybrid phenotype of macrophages and endothelial cells, which may regulate sinusoidal construction and Treg cell function. This study provides a comprehensive atlas that covers all hepatic cell types instrumental for further dissection of liver development, metabolic functions and diseases.

**In Brief:**

- Single cell transcriptomics of all hepatic cell types in neonatal and adult livers
- Concerted development of zonated metabolic functions in hepatocytes and NPCs
- Transient emergence of a distinct group of macrophages at postnatal day 7
- Hepatic cell-cell communications that program postnatal liver development

## INTRODUCTION

The liver is the most important organ for metabolic homeostasis and executes vital functions, including metabolism, detoxification, bile secretion, and production of plasma proteins. Hepatocytes are the main parenchymal cells, accounting for 80% of the liver volume and carrying out most of the metabolic and synthetic functions (Miyajima et al., 2014). In adult liver, the basic architectural unit is the zonated liver lobule, in which portal vein (PV) and hepatic artery and bile duct form a portal triad in the corner, with the central vein (CV) in the lobule center. Blood enters the lobules from PV and hepatic artery, and exchanges oxygen and nutrients with hepatocytes, while flowing through sinusoids. This unique architecture creates gradients of oxygen, nutrients and hormones, a phenomenon known as metabolic zonation between periportal and perivenous areas (Jungermann and Kietzmann, 1996). Hepatocytes locating in different layers along the lobule axis express different receptors, translocators, enzymes, and thus exhibit functional heterogeneity between zones. Given that metabolic syndromes are viewed as the emerging disease of the 21^st^ century, together with the rapid increase of liver cancer incidence and mortality, elucidating how the complex architecture and functions are developed is essential for understanding the molecular mechanisms of liver diseases and cancer and for designing efficacious therapeutic strategies.

Liver development is initiated around E8.5–E9.0 in mouse embryos by outgrowth of the primary liver bud from the ventral wall of the foregut (Zaret, 2002; Zhao and Duncan, 2005). This process is driven by concerted activities of fibroblast growth factors (FGFs) from the adjacent cardiac mesoderm and bone morphogenetic proteins (BMPs) from septum transversum mesenchyme (STM). By E9.5, the parenchymal progenitor cells delaminate from the bud and invade the surrounding STM as cords of hepatoblasts that can differentiate into hepatocytes and cholangiocytes, the biliary epithelium. Subsequent liver organogenesis that generates the complex architecture involves extensive differentiation of parenchymal and non-parenchymal cell (NPC) types, development of the biliary tract, sinusoidal capillaries and vasculature, and organization of extracellular matrix. Remarkably, the fetal liver is the main site of hematopoiesis, harboring hematopoietic stem cells, from E10.5 to E16.5 (Crawford et al., 2010; Sasaki and Matsumura, 1986). During the late stage of embryogenesis, the liver transitions to a primary organ of metabolism, following migration of hematopoietic stem cells to bone marrow (Gordillo et al., 2015; Zaret, 2002).

Single cell RNA-sequencing (scRNA-seq) has been used to characterize the landscape of the mouse gut endoderm and gastrulation (Nowotschin et al., 2019; Pijuan-Sala et al., 2019), and to dissect differentiation of the bipotential hepatoblasts into hepatocyte and cholangiocyte lineages (Yang et al., 2017). A robust and comprehensive scRNA-seq analysis was performed to characterize the onset of liver development from specification of endoderm progenitors to parenchymal and non-parenchymal cell lineage diversification and establishment in E7.5-E10.5 mouse embryos (Lotto et al., 2020). Another study interrogated single cell transcriptomes in endodermal and hepatic cells from E7.5 to E15.5 in Foxa2^eGFP^ mouse line (Mu et al., 2020).

Although a wealth of knowledge has been obtained regarding the initial stages of hepatic cell origin, specification and differentiation in embryos, much less is known about the neonatal and postnatal stages of liver development and maturation. At postnatal stage, hepatocytes organize into hepatic lobules, establishing the zonation structure. However, the driving factors of zonation construction and how hepatocytes adopt specific metabolic functions remains to be determined. One microarray analysis demonstrated four developmental stages from E11.5 to normal adult liver with distinct transcriptome profiles (Li et al., 2009). The dividing points of four stages are E14.5, E17.5 and Day 3. Another study using bulk RNA sequencing focused more on postnatal development from E17.5 to day 60 (Gunewardena et al., 2015). However, the bulk data analyses provided only the average values of different cell types, thus confounding the critical information on changes of cell type composition, cell-cell interactions and functional heterogeneity. Combined use of scRNA-seq with single-molecule fluorescence *in situ* hybridization allowed reconstruction of liver zonation and inference of lobule coordinates in fasted adult mouse, with over half of the liver genes shown to be significantly zonated, albeit the analysis was focused on hepatocytes only (Halpern et al., 2017).

In this study, we were able to capture all major hepatic cell types for comprehensive analysis of progressive liver development in mice after birth at single cell resolution. We characterized the process of hepatocytes and sinusoidal endothelial cell zonation establishment, and identified upstream regulators in hepatocyte development and their putative roles in tumorigenesis. We delineated development of the circadian rhythm in the liver as well as response of several critical metabolic pathways in hepatocytes after birth, including glycolysis, fatty acid β-oxidation, fatty acid and cholesterol biosynthesis. Remarkably, we identified a special group of macrophages, enriched on postnatal day 7, and inferred their role in regulating sinusoidal vascularization and regulatory T (Treg) cell activity via cell-cell interaction analysis. Herein we present a detailed atlas on postnatal liver development and maturation at single cell resolution.

## RESULTS

### scRNA-seq identifies distinct hepatocyte subpopulations during postnatal development

To investigate liver development, function and diseases at single cell resolution, we established a protocol for efficient isolation of all hepatic cell types, including hepatocytes and non-parenchymal cells (NPC), at high quality and integrity from mouse liver (see Methods). With this new protocol, our first aim was to interrogate how the liver develops and matures into the major metabolic organ after birth. For this purpose, we collected hepatic cells at postnatal day 1, 3, 7, 21, and 56 (marked as D1, D3, D7, D21, and D56), which covered the whole period of liver development from newborn to adult. All isolated single cells were used for scRNA-seq on 10x Genomics platform. Of a total of 65,891 cells subjected to RNA sequencing, 52,834 cells passed quality control based on the number of genes expressed, the count of raw reads and the mitochondrial gene percentage. Cells from all five time points were normalized and scaled for clustering and dimensional reduction using UMAP (Figures 1A, 1C). We identified a total of 31 cell types or cell states from all ages, based on expression of manually selected well-known markers (Figure 1B). Within the 31 clusters, we identified three groups of hepatocytes (Figure 1A). Hep-neonatal consisted of hepatocytes collected at the early stages, including D1, D3, and D7, while Hep-D21 and Hep-D56 included hepatocytes collected at D21 and D56, respectively.

**Figure 1.**
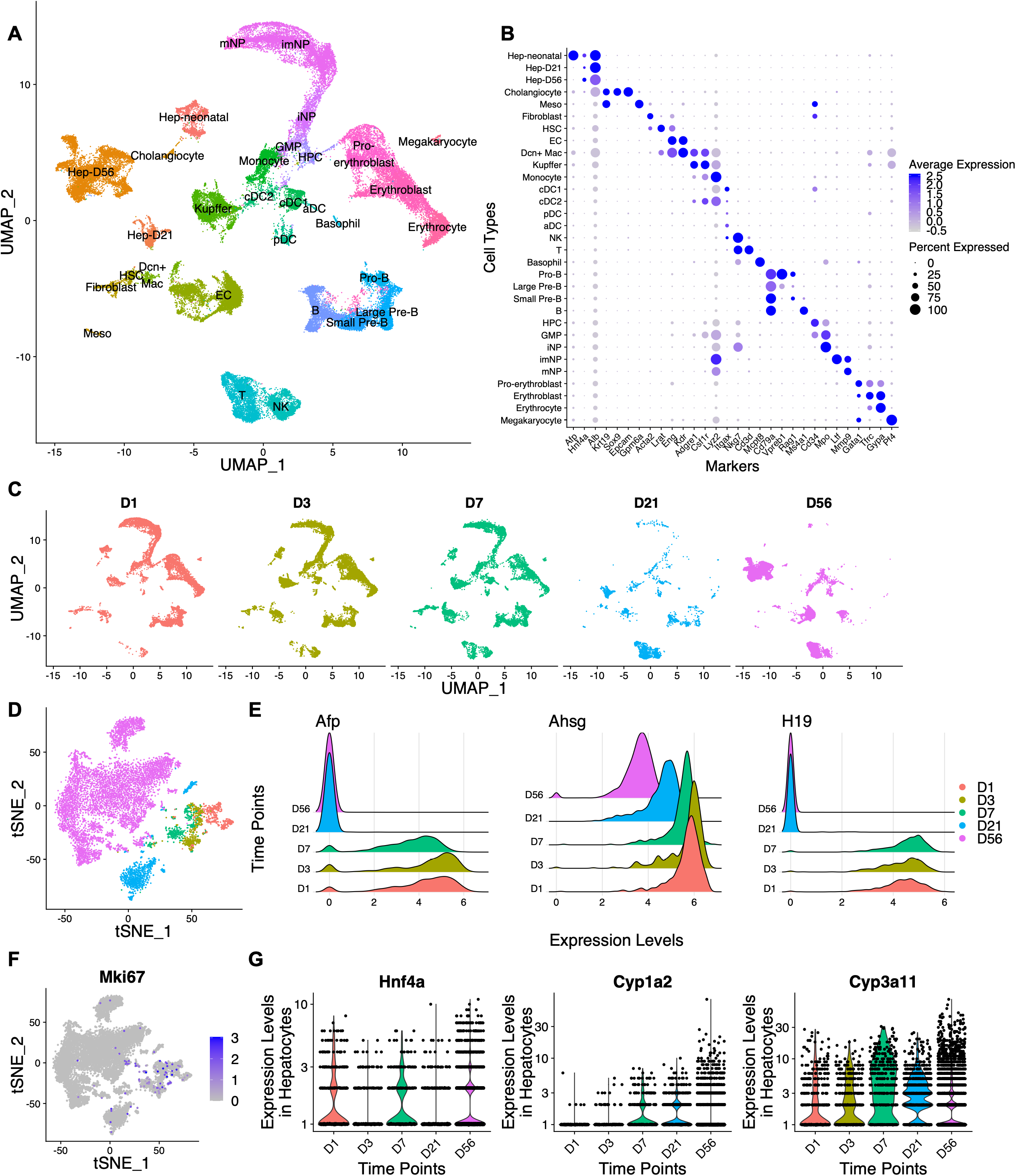
scRNA-seq identifies the major cell types in developing and adult liver. (A). UMAP visualization of liver cells from D1, D3, D7, D21 and D56. In total, 52,834 cells passed quality control. Colors indicate cell types, including Hepatocytes (Hep-neonatal from D1, D3 and D7; Hep-D21; Hep-D56), Endothelial cell (EC), Hepatic Stellate cell (HSC), Cholangiocyte, Fibroblast, Mesothelial cell (meso), Megakaryocyte, Erythroid cells (Pro-erythroblast, Erythroblast, Erythrocyte), T cell, Natural Killer (NK) cell, B cells (Pro-B, Large Pre-B, Small Pre-B, B), Dendritic cells (classical dendritic cell 1 – cDC1, classical dendritic cell 2 - cDC2, Plasmacytoid dendritic cell - pDC, activating dendritic cell - aDC), monocyte, Dcn^+^ macrophage (Dcn^+^ Mac), Kupffer cell, Neutrophils (immature neutrophil – iNP, intermediate mature neutrophil – imNP, mature neutrophil - mNP), Basophil, Granulocyte-Monocyte Progenitor (GMP) and Hematopoietic Progenitor cell (HPC). (B). Expression of selected markers for cell types. (C). UMAP visualization of liver cells from D1, D3, D7, D21 and D56, respectively. Colors indicate time points (n = 2∼4 for each time point). (D). tSNE map of hepatocytes from D1, D3, D7, D21 and D56. Cells in Hep-neonatal, Hep-D21 and Hep-D56 were segregated and re-analyzed. (E). Ridge plots displaying the expression levels of indicated markers at five time points. (F). tSNE map displaying *Mki67* (encoding KI-67) expression in hepatocytes from Figure 1D. (G). Violin plots displaying the expression levels of indicated mature hepatocyte markers at five time points.

Next, we focused on 9137 assigned hepatocytes from D1 to D56, expressing well-known biomarkers *Alb* and *Hnf4a* (Figure 1B). As expected, these cells were clustered primarily by time point (Figure 1D), with hepatocytes from D1, D3 and D7 locating close to each other, relative to those from later stages of D21 and D56. Of note, a few hepatoblast or hepatocellular carcinoma (HCC) markers, such as *Afp*, *Ahsg* and *H19*, were highly expressed by hepatocytes at three neonatal time points (Figure 1E), indicating that these cells are not fully differentiated mature hepatocytes. Indeed, hepatocytes from D1, D3 and D7 were actively proliferating as evidenced by the high expression of *Mki67*, encoding the cell proliferation marker Ki67 (Figure 1F). Hepatocytes from D21 represented a unique group of hepatocytes at an intermediate time point. These hepatocytes were not actively proliferating, with low expression of *Mki67* (Figure 1F); they also expressed low levels of *Afp*, *Ahsg* and *H19*, which were high in neonatal liver (Figure 1E). However, they were not fully matured either, expressing relatively low levels of mature hepatocyte markers, including *Hnf4a*, *Cyp1a2*, and *Cyp3a11* (Figure 1G), thus distinguishing them from hepatocytes in adult liver (D56).

We further investigated the dynamic changes of hepatocyte subpopulations emerging during postnatal development and identified a total of eight subpopulations, labelled as Hep1 to Hep8. The subpopulations with similar transcriptomes across multiple time points were given the same name. At each time point, we were able to identify two or three subpopulations (Figures 2A-2E). At neonatal stage, Hep1 and Hep2 were shared by D1, D3 and D7, while Hep3 and Hep4 were only identified at D1 and D7, respectively (Figures 2A-2C). Cells in Hep1 showed high expression of ribosomal proteins, indicating extremely active protein translation activity; Cells in Hep2 expressed typical liver metabolism-related genes, including *Apoa5*, *Apob*, and *Cyp4a14* (Figure S1A). At later stages, D21 and D56 livers shared two common rare subpopulations, Hep6 and Hep7, but had distinct major subpopulations, Hep5 and Hep8, respectively (Figures 2D-2E). Of note, the Hep3 subpopulation, only identified at D1, was featured by unique expression of *Scd2* (Figures 2F, S1A), which encodes one of the four mouse isoforms of stearoyl-CoA desaturase (SCD), a key enzyme for biosynthesis of monounsaturated fatty acids (Enoch et al., 1976; Miyazaki et al., 2005; Miyazaki et al., 2003). Among the four isoforms, SCD1 is the only one identified in adult liver, and is known to play a critical role in lipid metabolism. A previous report showed SCD involvement in Wnt signaling and its expression in activated hepatic stellate cells (HSC) and isolated tumor initiating cells (Lai et al., 2017). However, much less is known about SCD2 relative to SCD1. We checked *Scd1* and *Scd2* expression in hepatocytes across all time points (Figures 2G-2J). *Scd1* was mainly identified at D21 and D56, with highest expression at D56, while *Scd2* was mainly expressed in Hep3 at D1. Therefore, young and adult hepatocytes express two different SCD isoforms, with *Scd2* being an important gene in the neonatal liver and *Scd1* gradually becoming dominant in mature liver. An age-related expression pattern of *Scd1* and *Scd2* was also found in HSC (Figures S1B-S1C) and Kupffer cells (Figures S1D-S1E). Unlike hepatocytes, *Scd1* expression peaked at D21 in HSC and Kupffer cells, which might indicate a compensatory mechanism in these two cell types for low *Scd1* expression in hepatocytes at this stage, relative to adult liver. Interestingly, the expression of *Scd2*, but not *Scd1*, was elevated in Myc-induced tumor cells (unpublished dataset; Figures S1F-S1I), consistent with the previous *in vitro* data of isolated tumor cells (Lai et al., 2017).

**Figure 2.**
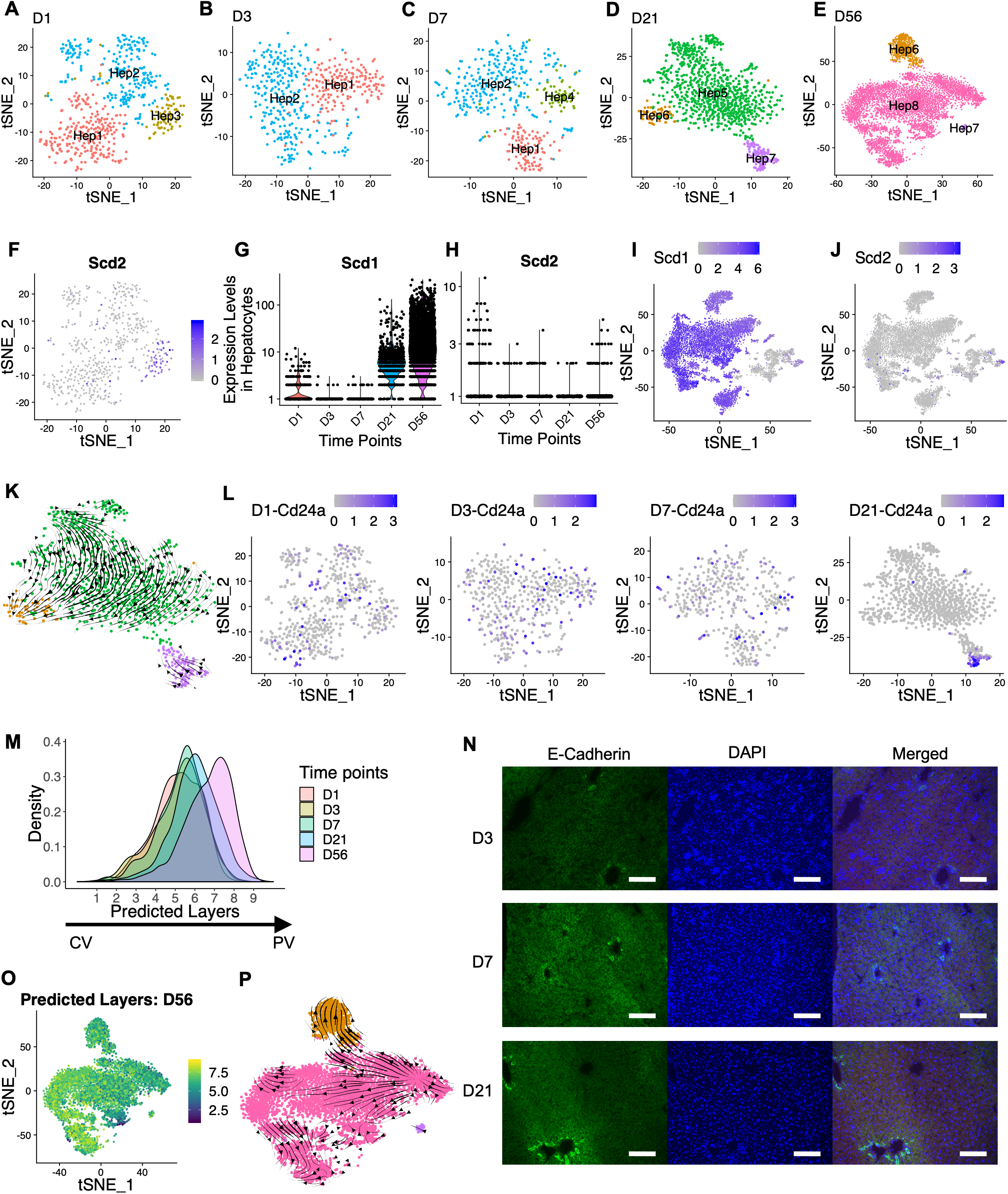
scRNA-seq identifies distinct transcriptome profiles in hepatocytes at each time point. (A-E). tSNE map of hepatocytes from D1 (A), D3 (B), D7 (C), D21 (D) and D56 (E). (F). tSNE map displaying *Scd2* expression in D1 hepatocytes. (G-H). Violin plots displaying *Scd1* (G) and *Scd2* (H) expression in hepatocytes from D1, D3, D7, D21 and D56. (I-J). tSNE map displaying *Scd1* (I) and *Scd2* (J) expression in hepatocytes from D1, D3, D7, D21 and D56. (K). RNA velocities visualized on tSNE map from Figure 2D (D21). (L). tSNE map displaying expression of *Cd24a* in hepatocyte at indicated time point. (M). Density plot displaying distribution of predicted layers of hepatocytes from D1, D3, D7, D21 and D56. Layer 1 represents peri-central hepatocyte; Layer 9 represents peri-portal hepatocyte. (N). Immunostaining of E-Cadherin in D3, D7 and D21 livers. Scale bar, 100 *µ*m. (O). Predicted layers visualized on tSNE map from Figure 2E (D56). (P). RNA velocities visualized on tSNE map from Figure 2E (D56).

We also performed RNA velocity analysis (Bergen et al., 2020; La Manno et al., 2018), to predict developmental trajectories at each time point. Anchoring at Hep5 as the most abundant subpopulation at D21, we identified clear trajectories showing that Hep5 originated from Hep7 and proceeded to Hep6, a more mature state (Figure 2K). A similar velocity pattern was also found among the equivalent subpopulations at D56 (Figure 2P). Cells in Hep7 highly expressed gene *Cd24a* (Figure S1J), which was proposed as a surface marker of hepatoblasts (Ochsner et al., 2007) and hepatocyte progenitor cells in adult mouse liver (Qiu et al., 2011). This gene was evenly expressed by hepatocytes in all subpopulations at neonatal stages, while its expression was constrained within Hep7 at D21 (Figure 2L), consistent with a differentiation potential of Hep7 cells revealed by RNA velocity analysis. Hep6 cells, the end point of D21 developmental trajectory, displayed high expression of *Neat1*, *Malat1* and *Mlxipl* (Figures S1J-S1K). Interestingly, *Neat1* and *Malat1* are well-known long non-coding RNAs (lncRNAs) implicated in malignant diseases, including liver and breast cancer (Malakar et al., 2017; Shin et al., 2019; Zhang et al., 2017). A similar cluster of cells was identified in human HCC patient samples (Sharma et al., 2020) and in our Myc-induced HCC mouse model (unpublished data), with co-expression of *Neat1*, *Malat1* and *Mlxipl*. Together, these results suggest that the Hep6 subpopulation identified at D21 and D56 may represent a special state of hepatocytes implicated in liver development and tumorigenesis. Besides their reported role in tumorigenesis, *Neat1* and *Malat1* represent the most abundant lncRNAs in liver nuclei. *Mlxipl* is one of the few predominantly spliced polyadenylated protein-coding mRNAs showing nuclear retention phenomenon (Bahar Halpern et al., 2015), which increases cytoplasmic mRNA stability against gene expression fluctuations. Thus, the RNA velocity analysis suggests that Hep6 is a stable and well-differentiated subpopulation in the liver.

We next explored how hepatocytes distributed spatially within the liver zonation structure across all time points. First, we checked expression patterns of four zonation markers, *Cyp2f2*, *Cyp2e1*, *Cdh1*, and *Glu1* (Figure S1L; (Braeuning et al., 2006). *Cyp2e1* was most abundant in the CV area and gradually decreased as moving toward the PV region, with *Cyp2f2* having an opposite pattern to *Cyp2e1*. *Glul*, encoding glutamine synthetase, was only expressed in the layer of hepatocytes around CV, while *Cdh1*, coding for E-Cadherin, was expressed in hepatocytes surrounding PV. The D56 hepatocytes were clearly distinguished based on their spatial locations, as shown by differential expression of these four zonation markers, which were not observed at any earlier time points. Previously, a combination of scRNAseq and smFISH were used to reconstruct mouse liver zonation profiles, by dividing hepatic lobules into nine layers and assigning individual cells to a layer based on the expression of landmark genes (Halpern et al., 2017). To investigate the zonation construction over developmental stages, we trained that published dataset (Halpern et al., 2017) with assigned layers and then predicted a spatial location of single cells from hepatocyte clusters at each time point (see Methods). Layer 1 represented hepatocytes in the CV area, while layer 9 was closest to the PV region. The dataset normalization, integration method and training model were selected with the smallest mean square error (MSE). The predicted layer distribution at D56 (Figure 2M) was consistent with the probabilities calculated previously based on the area of each layer in an ideal hexagonal lobule (Halpern et al., 2017), a model proposed for adult liver zonation, with the density of predicted layer peaking at layer 8. Interestingly, the predicted zonation profiles before D56 did not show similar patterns (Figure 2M), indicating that the zonation profile is progressively built up during postnatal liver development. The percentages of periportal hepatocytes gradually increased from D1 to D56, reflecting the construction process of the liver architecture, especially the portal triads, before liver maturation. This conclusion was further supported by immunostaining of E-Cadherin, which showed steadily increasing expression in periportal hepatocytes as a hallmark from D3 to D21 (Figure 2N). We then mapped the predicted layers back to tSNE visualization at D21 and D56 (Figures 2O, S1M). Although we identified a layer of E-Cadherin-positive hepatocytes surrounding portal vein at D21 (Figure 2N), transcriptome profiling showed that most hepatocytes from different layers were still mixed together (Figure S1M), indicating that D21 hepatocytes do not yet share different metabolic labors as in adult liver. Consistent with the expression of zonation markers (Figure S1L), hepatocytes could only be distinguished based on zonation at D56 (Figure 2O). Further, the combination of RNA velocity analysis (Figure 2P) and layer prediction (Figure 2O) at D56 indicated that the periportal hepatocytes were differentiated later, in agreement with the construction of periportal hepatocyte population from D1 to D56 (Figure 2M).

### Identification of transcription factors in programming liver development

We investigated the developing program of liver functions temporally, by focusing on the trajectory paths of hepatocytes across all time points. First, we established the developing trajectory and calculated the pseudotime for each single cell (Figures 3A-3B), using Monocle (Qiu et al., 2017; Trapnell et al., 2014). The D56 hepatocytes were divided into three groups, D56-p.c., D56-mid and D56-p.p., based on their predicted layers, and randomly down-sampled in order to be comparable to other time points. Since liver samples were collected for scRNA-seq from D1 to D56, the real time points could be used to evaluate accuracy of the pseudotime calculation (Figure 3C); this approach enhanced the reliability of the pseudo-temporal analysis. At D56, the pericentral hepatocytes (D56-p.c.) showed up first in the predicted trajectory followed by the mid-lobular hepatocytes (D56-mid), and then the periportal hepatocytes (D56-p.p.). This result indicates that the transcriptomic profile of hepatocytes in the developing liver was closer to that of pericentral than periportal hepatocytes in adult liver. In agreement with this concept, the zonation prediction also showed that the ratios of periportal hepatocytes gradually increased to D56 (Figure 2M), indicating that portal triads are being constructed later during liver maturation.

**Figure 3.**
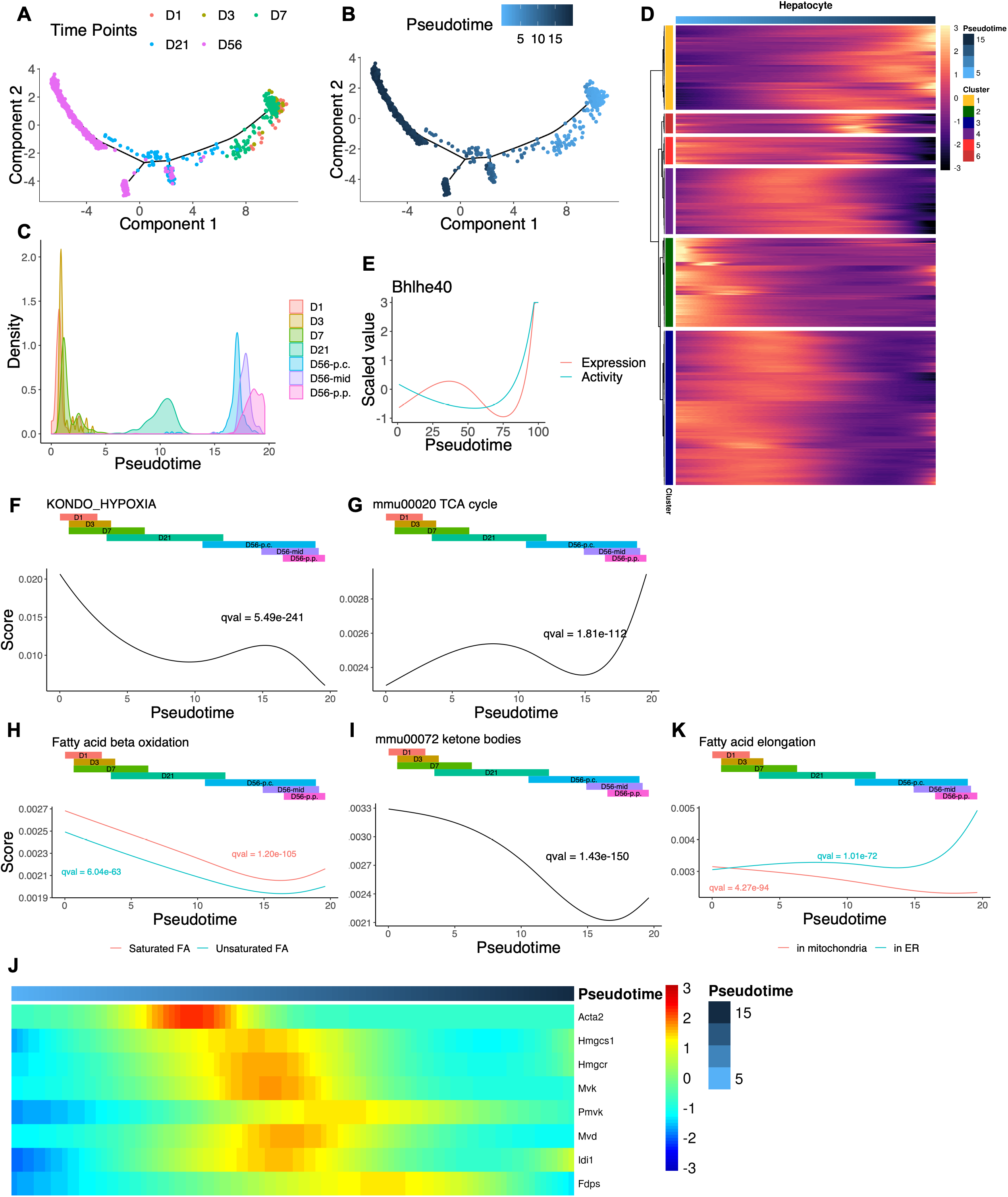
Dynamic changes of transcription factor activities and metabolic functions in hepatocytes. (A). Pseudotime analysis of hepatocyte development from D1 to D56 with Monocle 2. Colors indicate time points. (B). Pseudotime analysis from Figure 3A. Color indicates inferred pseudotime. (C). (C). Density plot displaying distribution of inferred pseudotime in Figure 3B. (D). Heatmap representing trends of differentially expressed genes as a function of inferred pseudotime in Figure 3B. (E). Fitted plot of Bhlhe40 scaled expression and activity values along pseudotime. The inferred pseudotime from Figure 3B was stretched from 0 to 100. (F-I). Fitted plots of enrichment scores for indicated pathways along inferred pseudotime in Figure 3B. (J). Heatmap displaying genes from the mevalonate pathway with differential expression along inferred pseudotime in Figure 3B. (K). Fitted plots of enrichment scores for indicated pathway along inferred pseudotime in Figure 3B.

We ordered single hepatocytes from all time points by pseudotime and performed differential expression analysis, to identify genes whose expression changed significantly as a function of pseudotime. These selected genes were then grouped into six clusters by pseudotemporal expression patterns (Figure 3D). For each cluster, we performed pathway enrichment analysis with simplified gene set collection from Molecular Signatures Databases (MSigDB) (Subramanian et al., 2005), with the significantly enriched pathways of each cluster listed in Table S1. The peripheral circadian clock-related pathway was identified in cluster 1, where genes were gradually increased from D1 to D56.

Furthermore, we searched for transcription factors (TFs) as possible regulators that control hepatocyte development. Accordingly, we performed single-cell regulatory network inference and clustering (SCENIC) (Aibar et al., 2017), to construct gene regulatory network (GRN) using the same subset of single cells for trajectory analysis. A TF activity matrix was generated with scores evaluating expression levels of downstream targets of each TF (regulon) in a single cell. Similar to the search for differentially expressed genes along pseudotime, we identified TFs with significantly changed activities as a function of pseudotime. This approach captured 36 TFs as possible candidates, showing significance on both activity and expression levels (Figures S2A-S2B). Included in the 36 candidates is Bhlhe40 (Dec1), a core TF involved in control of circadian rhythm (Honma et al., 2002; Sato et al., 2004). We fitted natural spline to visualize this TF’s change along pseudotime. Both its expression and activity increased dramatically at the late stage (Figure 3E), consistent with the pathway enrichment analysis results described above. Of note, we reported recently that the overall transcriptomic profiles in HCC were very similar to the young liver at the age of 1 month, reinforcing a notion of dedifferentiation during tumorigenesis (Wang et al., 2019). We asked if these identified regulators play important roles in HCC development. For each TF, we performed survival analysis of 371 patients’ data in the Liver Hepatocellular Carcinoma (LIHC) project from TCGA (https://www.cancer.gov/tcga). For TFs including *Klf9*, *Hbp1* and *Ets1*, whose expression and activity were low at early stage, patients with higher expression had better prognosis (Figures S2C-S2D). Conversely, for TFs such as *Tcf3* and *Hmgb2*, whose expression and activity were high at early stage, patients with higher expression showed worse prognosis (Figures S2C-S2D). Therefore, the liver development dataset constructed in this study may be instrumental for identification of candidate molecules as drivers and also therapeutic targets of HCC.

### Development of various metabolic functions in hepatocytes

Next, we explored development of metabolic functions in hepatocytes during the early postnatal stages. For each single cell, we calculated the pathway enrichment scores of gene sets collected from Kyoto Encyclopedia of Genes and Genomes (KEGG) database (Kanehisa and Goto, 2000) and MSigDB (Subramanian et al., 2005). We then ordered cells along pseudotime and filtered for gene sets with scores significantly changed along the trajectory. Furthermore, we visualized gene sets of interest through fitting a natural spline (see Methods) and also investigated the detailed changes of genes in a given gene set significantly expressed along the pseudotime.

In mammals, many cell types including hepatocytes use glycolysis as the main source of energy in the fetus due to low oxygen concentration and the immaturity of mitochondria. In the newborn, the metabolic energy source shifts as the activity of mitochondrial oxidative phosphorylation increases rapidly with air breathing (Böhme et al., 1983; Lindgren et al., 2019). As expected, we observed dramatic decrease in the expression of hypoxia-related pathway from D1 to D56 (Figure 3F). The hypoxia pathway was more prominently suppressed in the periportal zone than the pericentral zone at D56, with an opposite pattern observed for the tricarboxylic acid (TCA) cycle (Figure 3G). As glycolysis shares many enzymes with gluconeogenesis, we checked expression of genes specific for glycolysis or gluconeogenesis, or the reversible steps in the two opposing pathways (Figure S3A). The expression of most glycolysis-specific genes slowly increased when the glucose level restored and decreased subsequently, while gluconeogenesis-specific genes, including *Pcx* and *Fbp1*, were most highly expressed right after birth. This observation was consistent with sudden hypoglycemia at birth due to disruption of glucose supply from mother (Böhme et al., 1983), which could be remedied by enhanced gluconeogenesis. Similarly, glycogenolysis-specific genes also peaked at early stages (Figure S3B), to replenish glucose in blood, while glycogenesis showed up later and peaked in periportal hepatocytes at D56.

In addition to glucose metabolism, we observed a peak of β-oxidation of fatty acids immediately after birth, followed by a rapid decrease over time and slight increase in periportal hepatocytes at D56 (Figure 3H). This pattern indicates an essential role of fatty acid β-oxidation at early time points, producing large amounts of acetyl-CoA. As a downstream product of glycolysis, acetyl-CoA suppresses glycolysis while promoting gluconeogenesis to generate more glucose. Meanwhile, acetyl-CoA can be used to generate ketone bodies, because cells utilize ketones at the shortage of glucose (Beath, 2003). Indeed, ketone body-related pathway showed a similar pattern as β-oxidation, with a peak right after birth (Figure 3I). Besides its role in ketone production, acetyl-CoA is also a substrate of cholesterol biosynthesis. Therefore, we also checked critical genes involved in mevalonate pathway (Figure 3J). However, these genes were not highly expressed in newborns, but reached a peak at D21, showing that mevalonate pathway was not yet well established in neonatal livers.

Interestingly, fatty acid elongation in mitochondria and endoplasmic reticulum (ER) showed opposite patterns (Figure 3K). Fatty acid elongation happens in 1) cytosol, where the elongation is part of de novo fatty acid biosynthesis; 2) ER, the main site elongating 16-carbons or longer fatty acids in eukaryotic cells; and 3) mitochondria, a minor site elongating shorter fatty acids with 4-16 carbons (Jump, 2009). The ER-related gene set (GO_FATTY_ACID_ELONGATION_SATURATED_FATTY_ACID) includes most common fatty acid elongase subtypes, the rate-limiting enzymes for fatty acid elongation in ER. Our results show that short fatty acid elongation in mitochondria might be important for hepatocytes in newborn liver, while ER dominated this process later during development.

Several other interesting pathways were built up along developmental trajectory after birth and significantly zonated at D56. Bile secretion (Figure S3C), one of the major liver functions, peaked in the PV region at D56. Urea production from ammonia via major enzymes including CPS1, OTC, ASS, ASL and arginase (ARG1) also increased after birth and occurred in the PV region at D56 (Figures S3D). However, metabolism of xenobiotics showed an opposing pattern (Figure S3E). The pericentral hepatocytes more actively expressed xenobiotics metabolism related genes, which then decreased in the mid-lobe and PV regions.

### Development of liver endothelial and mesenchymal cells

Liver NPCs, which play important supportive and regulatory roles, include endothelial cells, HSCs, mesothelial cells, innate and adaptive immune cells. The endothelial cells are a large group of cells with high heterogeneity and multiple functions, which consist of liver sinusoidal endothelial cells (LSECs), macrovascular endothelial cells (MaVECs) and lymphatic endothelial cells. LSECs constitute the most abundant type of endothelial cells that form the wall of liver sinusoids. In adult liver, LSECs regulate hepatic blood flow, block the entry of blood pathogens, and also modulate functions of immune cells (Poisson et al., 2017). Liver endothelial cells across all five time points (D1-D56) were clustered into 11 subpopulations (Figure 4A), and non-LSECs shared similar transcriptomic profiles over the time period. LSEC at D56 constituted two large and continuous groups, periportal and pericentral LSECs (EC1, EC2) (Figures 4B, S4C). In contrast, we observed higher heterogeneity in pre-mature LSECs (EC3, EC4, EC5, EC6), without clear zonation separation. Furthermore, we identified EC9, a group of cells expressing proliferation and cell cycle related genes, which were detected at all time points (Figure 4B). All EC9 endothelial cells were in G2/M or S phase as predicted by cell cycle marker genes (Figure S4D), suggesting that a group of LSECs maintain the self-renewal capacity in developing and adult livers. MaVECs highly expressed a marker gene *Vwf*, without expression of the well-known LSEC markers, *Stab2* and *Lyve1* (Figure S4A), and these cells were further divided into two subpopulations based on their physical locations in liver. Central vein vascular endothelial cells (EC11, CVECs) expressed peri-central markers *Rspo3* and *Wnt2*, and portal vein vascular endothelial cells (EC7, PVECs) expressed peri-portal markers *Dll4* and *Efnb2* (Figure S4B; (Halpern et al., 2018). These markers effectively distinguish zonation for both LSECs and MaVECs. We also identified lymphatic ECs (EC10) expressing the marker gene *Thy1* (Figure S4A; (Jurisic et al., 2010). Interestingly, the LSEC markers *Stab2* and *Lyve1* were also detected in lymphatic ECs.

**Figure 4.**
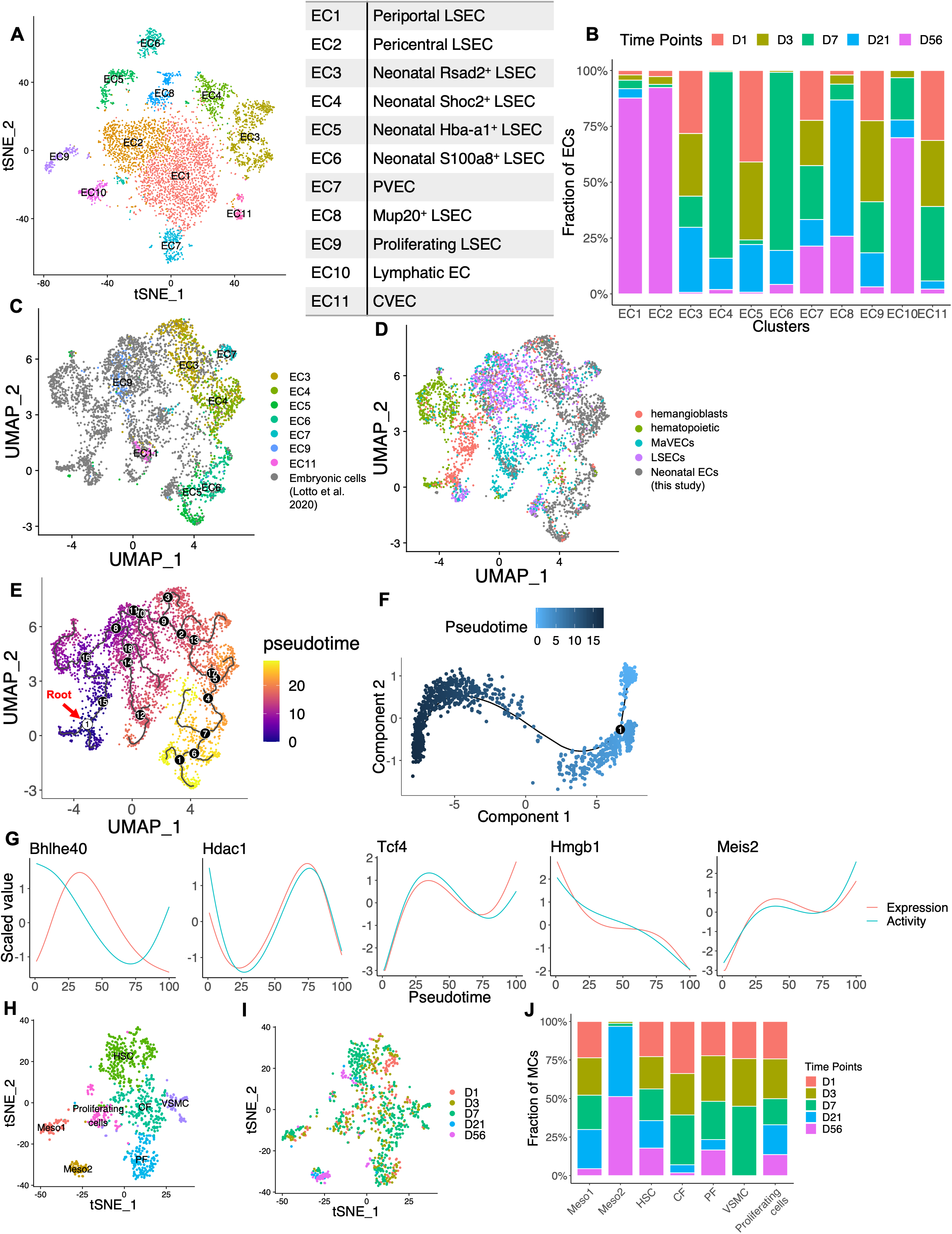
Development of liver endothelial cells and mesenchymal cells. (A). tSNE map of endothelial cells from D1, D3, D7, D21 and D56. Cells in EC were segregated and re-analyzed. Colors indicate assigned subpopulations. (B). Time point compositions of each endothelial cell subpopulation labelled in Figure 4A. (C). (C-D). UMAP visualization of cells from D1, D3, D7 and embryonic cells (including hemangioblasts, hematopoietic cells, HSEC and endothelium) from published data (Lotto et al., 2020). Color indicates endothelial cell subpopulation labelled in Figure 4A (C), or embryonic cell types (D). (E). Pseudotime analysis of cells from Figure 4C with Monocle 3. Color indicates inferred pseudotime. Root cell is labelled in white circle 1; branch points are labelled in black circles with numbers. (F). Pseudotime analysis of endothelial cells from D1, D3, D7, D21 and D56 with Monocle 2. Color indicates inferred pseudotime. (G). Fitted plot of scaled expression and activity values along pseudotime of indicated genes. The inferred pseudotime from Figure 4F was stretched from 0 to 100. (H). tSNE map of mesenchymal cells from D1, D3, D7, D21 and D56. Cells in meso, HSC and Fibroblast were segregated and re-analyzed. Colors indicate cell types. (I). tSNE map of cells from Figure 4H. Colors indicate time points. (J). Time point compositions of each cell type labelled in Figure 4H.

Previous single-cell analysis of embryonic liver from E7.5 to E10.5 (Lotto et al., 2020) showed a developmental trajectory from hemangioblasts to embryonic endothelial cells. To interrogate how the neonatal endothelial cells developed, we integrated their endothelial subset with our neonatal endothelial cells (Figures 4C-4D, S4E) and constructed a development trajectory from E7.5, E8.75, E9.5, E10.5, to D1, D3 and D7 (Figure 4E). The hemangioblast with highest differentiation potential, as calculated in the previous study, was defined as the root cell. Starting from the root, hemangioblasts differentiated into hematopoietic or endothelial cells at branch point 16. Later, at branch point 8, these early endothelial cells further differentiated into LSECs or MaVECs. Around this branch point, we noticed the emergence of proliferating LSECs after birth (EC9), which exhibited similar transcriptomic profiles as early embryonic endothelial cells. Within the branch of LSEC, neonatal cells developed in an order from EC3, EC4 to EC6 and EC5. Within the branch of embryonic MaVECs, we found the EC11 subpopulation, the neonatal central vein vascular ECs, showing a stable expression profile over time. In contrast, the EC7 subpopulation, consisting of neonatal portal vein vascular ECs, appeared later. These portal vein vascular cells possibly differentiated from EC3, the earliest subpopulation of neonatal LSECs. Taking together, our analysis showed that portal and central vein vascular endothelial cells used separate development paths. In particular, the portal vein area was constructed after birth, with concerted development of periportal hepatocytes and vascular endothelial cells.

Next, we investigated how neonatal LSECs developed into adult LSECs by pseudotemporal analysis of LSECs across the five time points. Similar to the methods described above for hepatocytes, we constructed a developmental trajectory and calculated pseudotime for each single cell (Figures 4F, S4F-S4G). This approach identified differentially expressed genes as a function of pseudotime, clustered by their pseudotemporal expression patterns (Figure S4H). We then performed pathway enrichment analysis for each cluster individually, with the significantly enriched pathways listed in Table S2. Among genes that decreased during postnatal development, we observed the enrichment of VEGFA-VEGFR signaling pathway, which may regulate endothelial cell proliferation and sinusoid construction at early time points. We also detected several glycolysis- and hypoxia-related pathways, showing that LSECs experienced similar metabolic changes as hepatocytes after birth. Meanwhile, *Vegf* expression can also be up-regulated under hypoxia by HIF-1 (Forsythe et al., 1996). For upregulated genes, there was a significant enrichment of interferon signaling pathways. Further, we identified TFs including Bhlhe40, Hdac1, Tcf4, Hmgb1 and Meis2, which exhibited significant expression and activity changes along LSEC developmental trajectory (Figure 4G). These TFs are possible regulators of LSEC maturation; among them, Meis2 was specifically expressed by endothelial cells in liver (Figure S4I). Surprisingly, Bhlhe40 showed opposite patterns of those identified in hepatocyte development (Figure 3E), with its role in LSECs remaining to be deciphered. However, it was reported that *Bhlhe40* expression was induced by hypoxia in vitro (Sato et al., 2008), and *Vegf* gene expression was regulated by circadian clock (Koyanagi et al., 2003). Thus, Bhlhe40 might contribute to the development and proliferation of LSECs in neonatal liver.

Another important group of liver NPCs is mesenchymal cells, which have been reported as the source of myofibroblasts after liver injury, especially HSCs (Iwaisako et al., 2014). We identified three main clusters of mesenchymal cells (Figure 1A), including HSCs, fibroblasts, and mesothelial cells (Meso). By segregating these cells for detailed analysis, we identified more mesenchymal cell types (Figure 4H) that were initially included within other mesenchymal cells. Interestingly, we detected two subgroups of mesothelial cells, Meso1 and Meso2, with distinct transcriptome profiles. Cells in both clusters highly expressed a mesothelial marker gene *Gpm6a* (Figure S4J). *Alcam*, a marker of embryonic mesothelial cells in mouse liver (Li et al., 2013), was expressed in Meso1, but not Meso2. Of note, Meso1 was mainly constituted by cells at the neonatal stage (Figures 4I-4J). A previous lineage tracing experiment showed that these mesothelial cells migrated inward from liver surface and differentiated into HSCs, fibroblasts, and smooth muscle cells during liver development (Li et al., 2013). Meso2 cells expressed *Igfbp4*, *Cd34* and *Clec3b*, and emerged at D21 and D56, exhibiting a gradual replacement of Meso1 by Meso2 during postnatal development. Additionally, we detected two fibroblast clusters, capsular fibroblast (CF) and portal fibroblast (PF), distinguished by their localization beneath the liver surface or near the portal triad, respectively (Balog et al., 2020). A group of proliferating cells with self-renewal capacity was also identified (Figure S4K), in which some cells expressed the HSC markers, *Reln* and *Lrat*, while others expressed the portal fibroblast marker, *Cd34* and *Clec3b* (Figure S4J).

### Dynamic changes of hematopoietic and immune cell populations

The major role of mammalian liver gradually transitions from hematopoiesis to metabolism around and after birth (Zaret, 2002). Our dataset from D1-D56 showed many clusters of developing hematopoietic cells, including hematopoietic progenitor cell (HPC), granulocyte-monocyte progenitor (GMP), as well as cells from B cell, neutrophil and erythroid lineages (Figure 1A). Both HPCs and GMPs expressed *Cd34*, a marker of hematopoietic precursors (Figure 1B). The development trajectories between these cells were inferred by force-directed graph (FDG) analysis (Figure 5A), which further verified some relationships shown in UMAP visualization (Figure 1A). HPCs were connected to three main paths, developing towards erythroid lineage, B cell lineage and GMPs, respectively. GMPs then gave rise to monocyte and neutrophil lineages. Based on markers used previously for bone marrow studies (Bjerregaard et al., 2003; Borregaard and Cowland, 1997), neutrophil lineages exhibited three major stages: 1) immature neutrophils (iNP; or neutrophilic promyelocytes), expressing high levels of *Mpo* and *Elane* (encoding elastase); 2) intermediate mature neutrophils (imNP; or neutrophilic myelocytes and metamyelocytes), expressing *Ltf* (encoding lactoferrin); and 3) mature neutrophils (mNP) (Figure 1B). Of note, this neutrophil lineage was not identified in human fetal liver in a recent scRNA-seq study (Popescu et al., 2019). Similarly, B cell lineage was also divided into several stages. First, the immunoglobulin heavy chain gene arrangement happened in pro-B cell, with a surrogate light chain expressed. This process required expression of *Rag1/2* for gene rearrangement and VpreB for surrogate light chain (Figure 1B). A real light chain rearrangement occurred in small pre-B cells expressing *Rag1/2*. Gene rearrangement, as well as the expression of *Rag1/2*, stopped in a middle stage, called large pre-B cells. We also divided the erythroid lineage into three stages based on reported markers (Elliott and Sinclair, 2012). These three lineages identified in postnatal liver shared similar stages of development to hematopoiesis observed in adult bone barrow. The percentages of developing hematopoietic cell types (Figures 5B, S5A-S5D) showed that the hematopoietic processes remained in the liver immediately after birth but receded rapidly, with most disappearing after D7. Development of T cell and NK cell lineages was not observed. It was reported previously (Rugh, 1990) that T cell development in liver occurred much earlier than B cells, with T cell progenitors migrating to thymus at about E13. The percentages of T cells and NK cells, as well as dendritic cells (DCs), gradually increased over time (Figures S5E-S5F), indicating their hepatic migration from thymus or bone marrow after birth.

**Figure 5.**
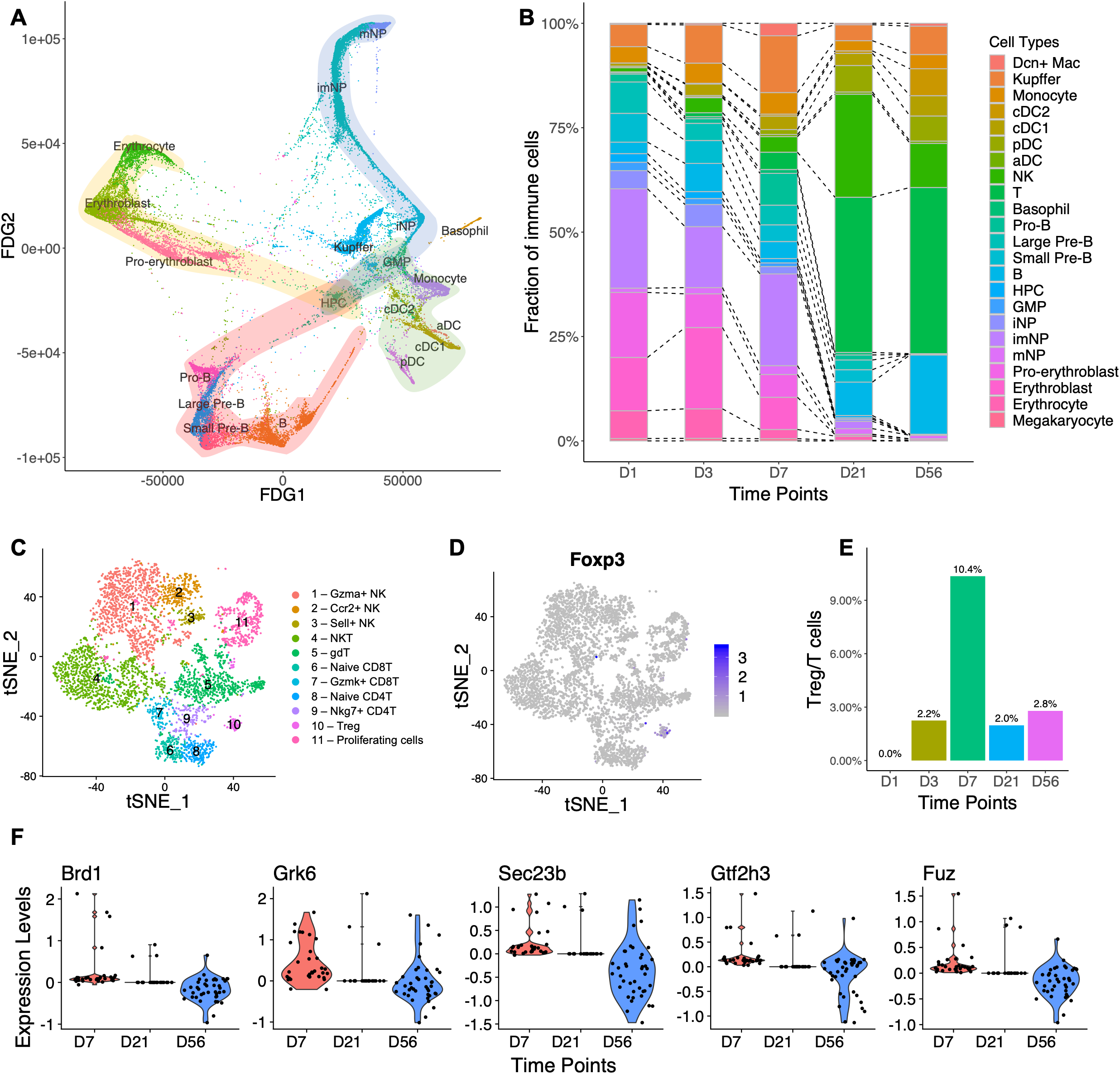
Changes of immune cell profiles during liver postnatal development. (A). Force-directed graph (FDG) of HPCs, GMPs, erythroid cells, neutrophils, B cells, DCs, basophils, monocytes and Kupffer cells. (B). Cell type compositions of immune cells at each time point. (C). tSNE map of T and NK cells from D1, D3, D7, D21 and D56. Cells in T and NK clusters were segregated and re-analyzed. Colors indicate cell types. (D). tSNE map displaying Foxp3 expression in cells from Figure 5C. (E). Percentages of Treg cells out of all T cells at indicated time point. (F). Violin plots displaying differentially expressed genes in Treg cells at D7, compared to Treg cells at D21 and D56. D1 and D3 were not included because of too few Treg cell numbers.

Since T cells and NK cells are highly heterogeneous, the observed percentage changes in T or NK cells from D1 to D56 (Figure S5E) might not depict the patterns for individual subtypes. Accordingly, we segregated these cells and analyzed them separately. Within the eleven T and NK cell subtypes identified (Figures 5C, S5G), Treg cells (Figure 5D) showed a distinctive percentage change over time (Figures 5E, S5H), with a peak at D7 relative to other time points. Consistent with this observation, a recent report (Li et al., 2020) showed accumulation of Treg cells in liver between D7 and D14 after birth. Here, we compared differentially expressed genes in Treg cells between D7, D21 and D56. Among the top-ranked genes (Figure 5F), *Brd1* and *Gtf2h3* were reported to be involved in cell proliferation or cell cycle progression (Drapkin et al., 1996; Mishima et al., 2011; Mishima et al., 2014). Tregs also highly expressed Grk6, which encodes a G-protein-coupled receptor kinase responsible for CXCR4 phosphorylation following CXCL12 treatment, and leading to immune cell recruitment and enhanced immune functions (Busillo et al., 2010).

### A subpopulation of macrophages emerges transiently around postnatal day 7

One unique property of the liver is its strong innate immunity and possession of large numbers of macrophages, with Kupffer cells being the liver resident macrophages. We identified a Kupffer cell cluster, which localized close to dendritic cells (DCs) and monocytes on UMAP visualization (Figure 1A). This cluster of cells expressed high levels of the Kupffer cell markers *Adgre1* (encoding F4/80) and *Csf1r* (Figures 1B, 6A-6B), and the percentage of Kupffer cells peaked at day 7 during the time period examined (Figure 6D). We identified six subpopulations, KC1-KC6, when analyzing Kupffer cells separately (Figures 6C, S6A-S6B), with distinct gene expression profiles (Figure S6C) and TF activities (Figure S6D). The KC6 group emerged at D1, with high expression of *Ccl9* and high TF activity of Nfil3. KC5 mainly came from D56, with high expression of *Ly6a* and *Cxcl13*, as well as high TF activity of Zbtb7a and Irf2. Other subpopulations (KC1-KC4) consisted of Kupffer cells from D1 to D21, with most gene expression patterns conserved over time. Among them, KC1 showed high levels of protein translation, with highest expression of ribosomal proteins and Myc activity, and KC4 included a group of actively proliferating cells.

**Figure 6.**
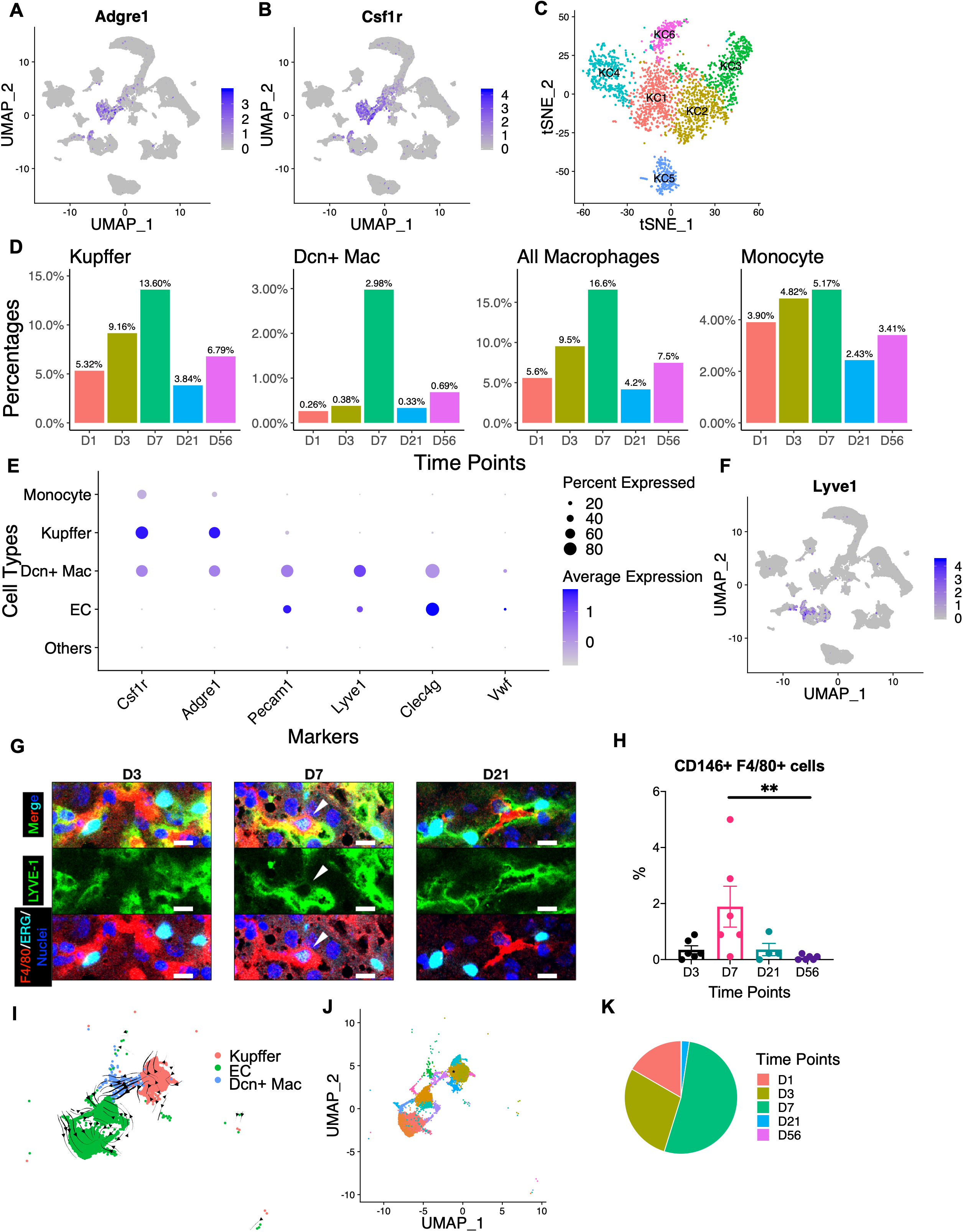
A unique subtype of macrophages identified at postnatal day 7. (A-B). UMAP visualization of *Adgre1* (A) and *Csf1r* (B) expression levels in all cells included in this analysis (Figure 1A). (C). tSNE map of Kupffer cells from D1, D3, D7, D21 and D56. Cells in Kupffer clusters were segregated and re-analyzed. Colors indicate assigned subpopulations. (D). Percentage of indicated cell types out of total immune cell population at each time point. (E). Dot plots displaying expression levels of selected markers for indicated cell types. (F). UMAP visualization of *Lyve1* expression levels in all cells included in this analysis. (G). Immunostaining of F4/80, ERG and LYVE-1. Representative image taken under confocal microscope. Dcn^+^ Mac cells were indicated by white arrowhead. Scale bar, 10 *µ*m. (H). Quantitative result of FACS analysis of CD146^+^ F4/80^+^ cells in isolated liver NPCs from D3, D7, D21 and D56, showing an enrichment at D7. Statistical analysis was done with Kruskal-Wallis test, followed by Dunn’s test. (** p < 0.01). (I). RNA velocities of Kupffer cell, EC and Dcn^+^ Mac from D1, D3, D7, D21 and D56, visualized on UMAP. (J). Clustering analysis of cells from Figure 6I. The Kupffer cell cluster showing differentiation potential to Dcn^+^ Mac was labelled by asterisk (transitioning KC). (K). Time point composition of asterisk labelled cluster from Figure 6J.

In this study, we found a novel cluster of macrophages (Dcn^+^ Mac). Besides the traditional Kupffer cell markers, *Adgre1* and *Csf1r* (Figures 1B, 6A, 6B, 6E), the Dcn^+^ macrophages also expressed general endothelial cell markers *Pecam1*, *Eng*, *Kdr*, and LSEC-specific markers *Lyve1* and *Clec4g*, but were negative for a MaVEC marker *Vwf* (Figures 1B, 6E-6F). Surprisingly, almost all the Dcn^+^ macrophages were identified at D7 (Figure 6D), which prompted us to launch a thorough interrogation of this unique cell group. First, we validated the existence of Dcn^+^ Mac and its abundance at D7 by co-immunostaining for the LSEC marker LYVE-1, the endothelial cell-specific TF ERG, and the macrophage marker F4/80. To confirm the co-expression of these markers in a single cell, instead of two neighboring cells, we included markers expressed on the endothelial cell surface or in the nucleus. Under a confocal microscope at higher resolution, we indeed visualized single cells co-expressing LYVE-1, ERG and F4/80 (Figure 6G). FACS analysis validated that this cell type emerged mainly at postnatal D7 (Figure 6H). Next, we investigated the possible origin of Dcn^+^ Mac cells. Given the expression of markers for both cell types, we reasoned that Dcn^+^ Mac might be either derived from LSEC and then gained macrophage’s transcriptomic profiles, or the converse. To address this question, we performed RNA velocity analysis for Dcn^+^ Mac, Kupffer and endothelial cells across all five time points. The result clearly showed a trajectory of Dcn^+^ Mac development from Kupffer cells rather than endothelial cells (Figure 6I). The group of Kupffer cells at the transition point (Figure 6J; transitioning KC, labelled by asterisk) were identified at D1, D3 and D7 (Figure 6K), showing a differentiation potential at the neonatal stage. Then, we mapped six Kupffer cell subpopulations in this UMAP (Figure S7A). We found that transitioning KC came primarily from KC2 and displayed high TF activity of Pou2f2 and Tcf7l2 (Figure S6D), which might drive the transition of Dcn^+^ Mac from KC2. Several recent reports (Chakarov et al., 2019; Lim et al., 2018) addressed LYVE1-expressing macrophages, which restrained tissue fibrosis in mouse lung and aorta, respectively. However, those cells were macrophages expressing only one endothelial cell marker, LYVE1. In contrast, the Dcn^+^ Mac detected in the liver at D7 had a fusion of macrophage and endothelial cell gene expression profiles. To interrogate which genes are significantly expressed in transitioning (asterisk-labelled) KC, we performed differential gene expression analysis between the transitioning KC and other Kupffer cells located far from Dcn^+^ Mac (Figure 6J). Surprisingly, most genes previously reported to be highly expressed in lung LYVE1-expressing macrophages including *Fcna*, *Marco*, *Cd163*, *Cd209f*, *Timd4* and *Mrc1* (Chakarov et al., 2019; Lim et al., 2018), were significantly expressed in this Kupffer cell population (Figure S7B).

We reasoned that this previously unrecognized Dcn^+^ Mac might be a unique cell type that appears transiently and plays a critical role during postnatal liver development. To test this conjecture, we performed ligand-receptor analysis using CellPhoneDB (Efremova et al., 2020) to identify interactions between Dcn^+^ Mac and other cell types at D7 and infer possible functions for Dcn^+^ Mac cells. We counted numbers of significant interactions between any two cell types (Figure 7A), with Dcn^+^ Mac as the communication hub. Cell types actively interacting with Dcn^+^ Mac included HSCs, fibroblasts, mesothelial cells, cholangiocytes, VSMCs and all subtypes of endothelial cells. Much fewer interactions were observed between these cell types and Kupffer cells (Figure 7B). We then examined specific signaling crosstalk related to Dcn^+^ Mac, as compared to Kupffer cells (Figure 7C). To explore possible influences of Dcn^+^ Mac on other cell types, we focused on interaction pairs with ligands secreted by Dcn^+^ Mac and receptors expressed on other cell types. Surprisingly, we detected several VEGF signaling-related pairs between Dcn^+^ Mac and LSEC, none of which were identified between Kupffer cells and LSEC, suggesting a role of Dcn^+^ Mac in LSEC proliferation, angiogenesis and vascularization. Of note, Dcn^+^ Mac also secreted CXCL12 which binds to CXCR3 and CXCR4 on Treg cells. Together with the activity of *Grk6*, which has elevated expression level in D7 Treg cells (Figure 5F), binding of CXCL12 has been demonstrated to significantly activate CXCR4 signaling (Busillo et al., 2010). Activation of this signaling axis contributed to Treg recruitment and enhanced Treg cell activity, and may thus explain the wave of Treg observed at D7 (Figure 5E).

**Figure 7.**
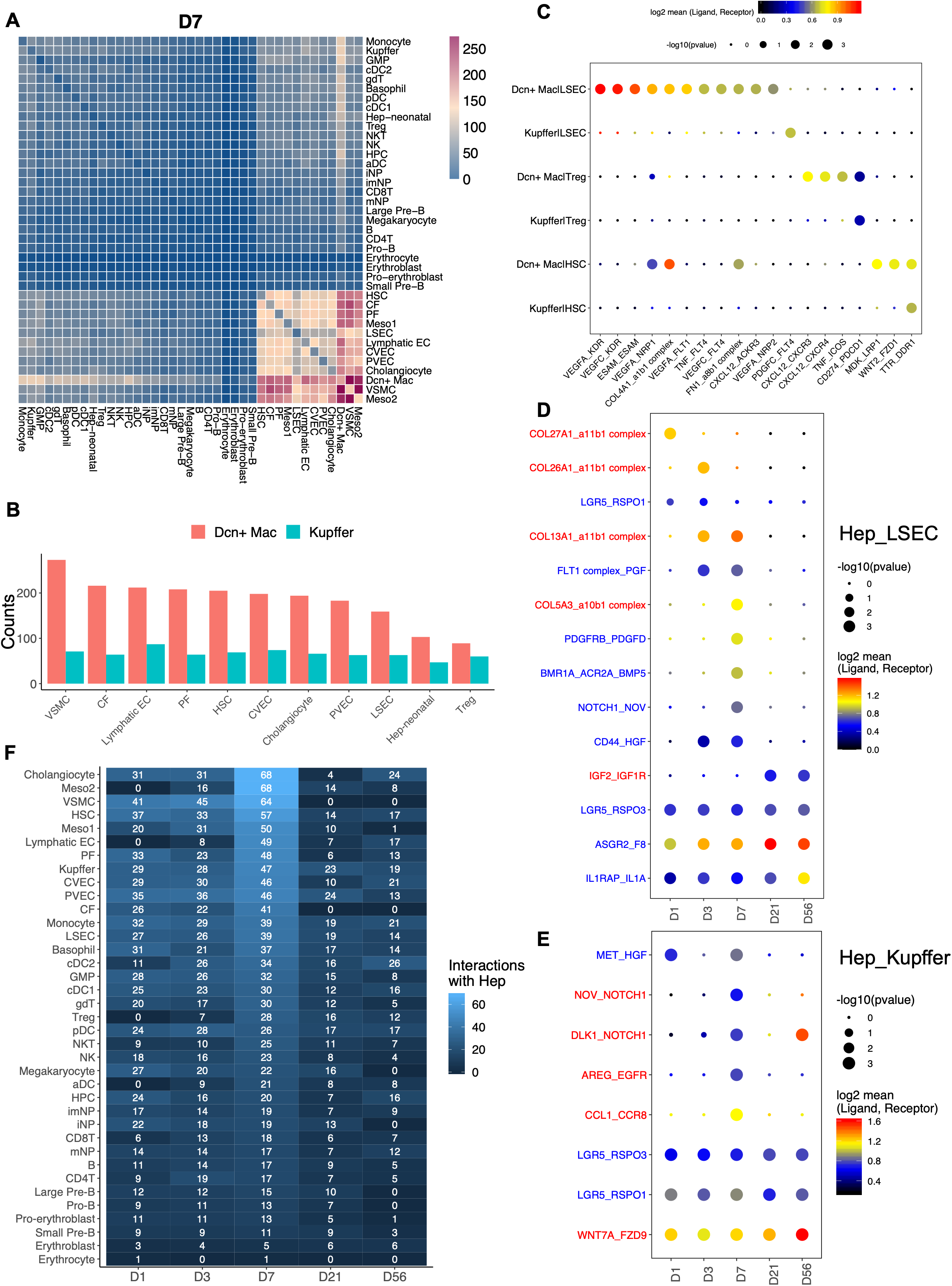
Predicted hepatic cell-cell interactions. (A). Heatmap displaying the total numbers of interactions between each pair of cell types at D7. (B). Bar plot comparing numbers of interactions between indicated cell types and Dcn^+^ Mac or Kupffer cells. (C). Dot plot of selected ligand-receptor interactions between Dcn^+^ Mac and indicated cell types (Dcn^+^ Mac|LSEC, Dcn^+^ Mac|Treg, Dcn^+^ Mac|HSC) or between Kupffer and LSEC (Kupffer|LSEC, Kupffer|Treg, Kupffer|HSC). Ligand and receptor were expressed by corresponding cell types. For example, ‘moleculeA_moleculeB in cellC|cellD’ indicates the putative interaction between molecule A expressed by cell type C and molecule B expressed by cell type D. P values are indicated by circle sizes, and the means of the average expression levels of interacting ligand and receptor are indicated by color. (D-E). Dot plot of selected ligand-receptor interactions between hepatocytes and LSEC (D) or Kupffer cells (E). Interacting pairs in red indicate ligands secreted by hepatocytes; interacting pairs in blue indicate ligands secreted by LSEC or Kupffer cells. (F). Heatmap displaying the total numbers of interactions between NPCs and hepatocytes at five time points.

### Hepatic cell-cell communications

Using the same method described above, we also checked crosstalk between hepatocytes and NPCs, and interrogated how these signaling events changed over the early postnatal stages. We performed ligand-receptor analysis for all five time points individually. To investigate the cell-cell interactions that maintain and control hepatocyte zonation, we filtered for significant pairs whose receptors were expressed on hepatocytes and were previously shown to be zonated. Of note, we identified RSPO1/3-LGR5 interactions between LSECs and hepatocytes (Figure 7D). RSPO3-LGR5 was significantly expressed across all time points, while RSPO1-LGR5 was only identified at neonatal stage. The binding of RSPO to LGR5, present on pericentral hepatocytes, leads to increased Wnt signaling, which modulates zonation pattern and metabolic pathways in pericentral area (Planas-Paz et al., 2016; Rocha et al., 2015; Torre et al., 2010). This signaling was also identified between Kupffer cells and hepatocytes across all time points (Figure 7E). Compared to the pericentral area, much less is known about the molecular signaling events that modulate periportal hepatocyte zonation. Using the same method, we identified ASGR2-F8 interaction between hepatocytes and LSECs. ASGR2 is a marker of periportal hepatocytes (Halpern et al., 2018). We then counted the interactions between hepatocytes and all the other NPCs (Figure 7F). Surprisingly, for almost all cell types closely interacting with hepatocytes, the most active time point was D7, compared to other time points. Some interaction pairs, including PDGFRB-PDGFD between hepatocytes and LSECs (Figure 7D) and NOV-NOTCH1 between hepatocyte and Kupffer cells (Figure 7E), were only identified at D7. These data reveal that that D7 is a critical time point for establishing many aspects of mature liver architecture and function during postnatal development.

## DISCUSSION

By establishing an efficient cell isolation protocol, we successfully captured all mouse liver cell types from newborns and adults, at multiple time points between postnatal D1 - D56, for scRNA-seq analysis. This study provides the first detailed blueprint that describes stepwise changes at single cell resolution to illustrate how a neonatal liver develops and matures into the major metabolic organ. Trajectory analysis revealed progressive and concerted development and functional maturation of hepatocytes and sinusoidal endothelial cells in the establishment of metabolic zonation. Complex interactions between hepatic cell types are apparently involved in promoting and coordinating programs in the early stages of postnatal liver development. Remarkably, we identified a special type of macrophage, which emerges at D7, that exhibits a hybrid phenotype of both macrophages and endothelial cells. Intercellular interaction analysis demonstrated that this group of cells might play a critical role in regulation of liver sinusoidal vascularization and Treg cell activity.

Among all the liver cell types examined, hepatocytes displayed high levels of variation or heterogeneity in the early developmental stages after birth. Unlike other cell types, hepatocytes were clearly separated by developmental stages on UMAP (Figure 1A). In neonatal liver, hepatocytes highly expressed hepatoblast or HCC markers, including *Afp*, *Ahsg* and *H19*. We also identified a unique *Scd2*^+^ hepatocyte subpopulation at D1, whose expression was gradually replaced by *Scd1* during postnatal development, suggesting a specific role of *Scd2* in neonatal liver metabolism. A rare *Cd24a*^+^ hepatocyte subpopulation was identified at D21 and D56, representing a candidate for hepatocyte progenitors based on RNA velocity analysis, in agreement with previous data (Qiu et al., 2011). By combining these hepatocyte subpopulations, pseudotemporal analysis enabled us to explore the developmental trajectory based on the samples collected from multiple time points. The unbiased analysis of upstream regulators and enriched pathways showed an establishment of liver circadian clock after birth regulated by factors, such as Bhlhe40 (Dec1), and also displayed modulation of several critical metabolic pathways in adaptation to dramatic environmental changes in neonatal life. As the major liver function switches from hematopoiesis to metabolism toward the end of embryogenesis, it is reasonable to speculate that various metabolic pathways develop and eventually mature in adult liver. Strong support to this conclusion was rendered by a previous study (Nakagaki et al., 2018), which measured expression of 25 selected metabolism-related genes in liver lysates using real-time PCR. Indeed, we found most of these pathways changed in an increasing trend, although a few metabolic pathways peaked in newborns and decreased later. Due to a sudden hypoglycemia environment occurring at birth due to disruption of glucose supply from mother, neonates had two strategies to overcome this crisis, by enhancing gluconeogenesis and glycogenolysis, to produce more glucose. Another mechanism is to enhance β-oxidation to generate large amounts of acetyl-CoA for synthesis of ketone bodies as energy source. As a downstream product of glycolysis, acetyl-CoA accumulation suppresses glycolysis and promotes gluconeogenesis. Besides, we also found fatty acid elongation in ER increased, while elongation of shorter fatty acid in mitochondria decreased. The later process almost disappeared in adult liver, indicating mitochondria might be a transient place in neonatal liver for fatty acid elongation, and ER soon replaces mitochondria as the major site.

Zonation construction is a most critical event during liver development into the primary metabolic organ. Our data analysis showed clearly that the metabolic zonation structure is not yet developed before weaning at D21. Although immunostaining showed an E-cadherin^+^ layer of hepatocytes surrounding PV at D21, but not at D3 and D7 (Figure 2N), the combined transcriptome profiles and zonation prediction results indicate that the D21 hepatocytes had not shared different metabolic labors yet as in adult liver. Furthermore, the maturation of periportal area appears delayed, compared to pericentral area, as revealed by zonation prediction (Figures 2M) and pseudotemporal analysis with hepatocytes across the five time points (Figure 3C). Similar patterns of developmental process was observed for endothelial cells revealed by pseudotemporal analysis with integrated embryonic (Lotto et al., 2020) and postnatal datasets (Figures 4C-4E). Consistent with previous data (Planas-Paz et al., 2016; Rocha et al., 2015), we identified RSPO1/3-LGR5 interactions between LSECs and hepatocytes, and also between Kupffer cells and hepatocytes (Figures 7D-7E), suggesting an important role in maintaining and controlling hepatocyte zonation. Furthermore, other ligands secreted by LSEC or Kupffer cells, including PDGF, HGF, PGF and BMP, might also regulate hepatocyte proliferation and zonation remodeling at the early stage.

In previous experiments, we compared the bulk transcriptomic data between HCC tumors and 1-month-old young liver tissue, and observed high levels of similarity (Wang et al., 2019). We believe the shared expression patterns between an actively developing liver and liver tumor tissue will be instrumental for dissection of mechanisms underlying hepatocarcinogenesis and also for search of new biomarkers in HCC initiation and progression. Indeed, a recent report (Sharma et al., 2020) described a shared group of macrophages between HCC patients, human fetal liver and mouse embryonic liver. In the current study, we found that a rare hepatocyte subpopulation specifically expressed *Scd2,* which was enriched only in the D1 liver. In a Myc-induced HCC model, we also detected highly elevated expression of *Scd2* in tumor cells accompanied by reduced *Scd1* expression, showing a similar pattern as D1 hepatocytes. Several upstream regulators identified in liver postnatal developmental trajectory showed correlation with HCC patients’ survival. An immature liver circadian clock program was observed in neonatal liver before maturation. Of note, disruption of the circadian clock induced spontaneous HCC development, as reported previously (Kettner et al., 2016).

One most interesting part of our observations is the drastic change of the hepatic microenvironment during postnatal development. Starting around E13 to E14, hematopoietic cells migrate from embryonic liver to thymus, spleen and bone morrow (Crawford et al., 2010), leaving space for hepatocytes to expand. We did observe the complete processes of erythrocyte, neutrophil and B cell lineage differentiation from progenitor cells in the D1, D3, and D7 datasets (Figure 5A). Among all the environmental changes observed in neonatal livers, we focused on events that happened at D7, as this date appeared to be a critical turning point, consistent with previous bulk RNA-seq data of the liver (Gunewardena et al., 2015; Li et al., 2009). Indeed, we identified Dcn^+^ Mac and Treg cells enriched at D7, as well as active interactions between hepatocytes and other cell types. It must be indicated that Dcn^+^ Mac was a previously unrecognized subtype of cells with a hybrid phenotype of macrophages and endothelial cells. We proved their existence and enrichment at D7 with confocal microscopy and FACS analysis. In particular, the co-expression of LYVE-1, ERG and F4/80 in one single cell visualized under confocal microscope clearly demonstrated that Dcn^+^ Mac is not a doublet mistakenly detected by scRNA-seq or FACS. The Dcn^+^ Mac cells were very likely derived from Kupffer cells and acquired endothelial cell markers, and they actively interacted with LSEC, HSC, fibroblast, mesothelial cell and Treg cells. In particular, we showed that the CXCL12-GRK6-CXCR4 signaling between Dcn^+^ Mac and Treg cells may promote Treg cell recruitment and their immune suppressive activity

While this work fills in a gap of knowledge on the postnatal stage of liver development between embryonic and adult livers at single cell resolution, several key results do provide fresh views on previously unappreciated high levels of hepatocyte heterogeneity in the developing liver after birth, the progressive construction of metabolic zonation in hepatocytes from pericentral to periportal regions, and the concerted development of hepatocytes and neighboring endothelial cells, HSC, and Kupffer cells. Further dissection of the newly identified Dcn^+^ Mac cell functions and their interactions with other hepatic cell types will contribute to better understanding of how a unique immune-tolerant microenvironment is developed temporally in premature liver and how it is tightly controlled in adult liver.

## ACKNOWLEDGMENTS

We thank our lab members for helpful discussion. Single cell RNA-seq was conducted at the IGM Genomics Center, UCSD. This work was supported by NIH grants (R01DK128320, R01CA236074 and R01CA239629) to G.S.F. Y.L. received David V. Goeddel Chancellor’s graduate fellowship, J.L. was supported by a postdoc fellowship from the international liver cancer association, K.K. was supported by a postdoc fellowship from Moores UCSD Cancer Center.

## AUTHOR CONTRIBUTIONS

Conceptualization: Y.L., and G.S.F.; Methodology: Y.L., K.Z., and G.S.F.; Software: Y.L.; Investigation: Y.L., K.K., J.L., and B.X.; Data Analysis and Interpretation: Y.L., X.S., K.Z., and G.S.F; Writing – Original Draft: Y.L., and G.S.F.; Writing - Reviewing and Editing: Y.L., K.K., J.L., X.S., K.Z., and G.S.F.; Supervision and Funding Acquisition: G.S.F.

## DECLARATION OF INTERESTS

The authors declare no competing interest.

## SUPPLEMENTAL INFORMATION

**Table S1. Enriched pathways of six gene clusters grouped by pseudotemporal expression patterns, related to Figure 3D.**

**Table S2. Genes and pathways significantly changed during LSEC postnatal development, related to Figure S4H.**

**Figure S1.**
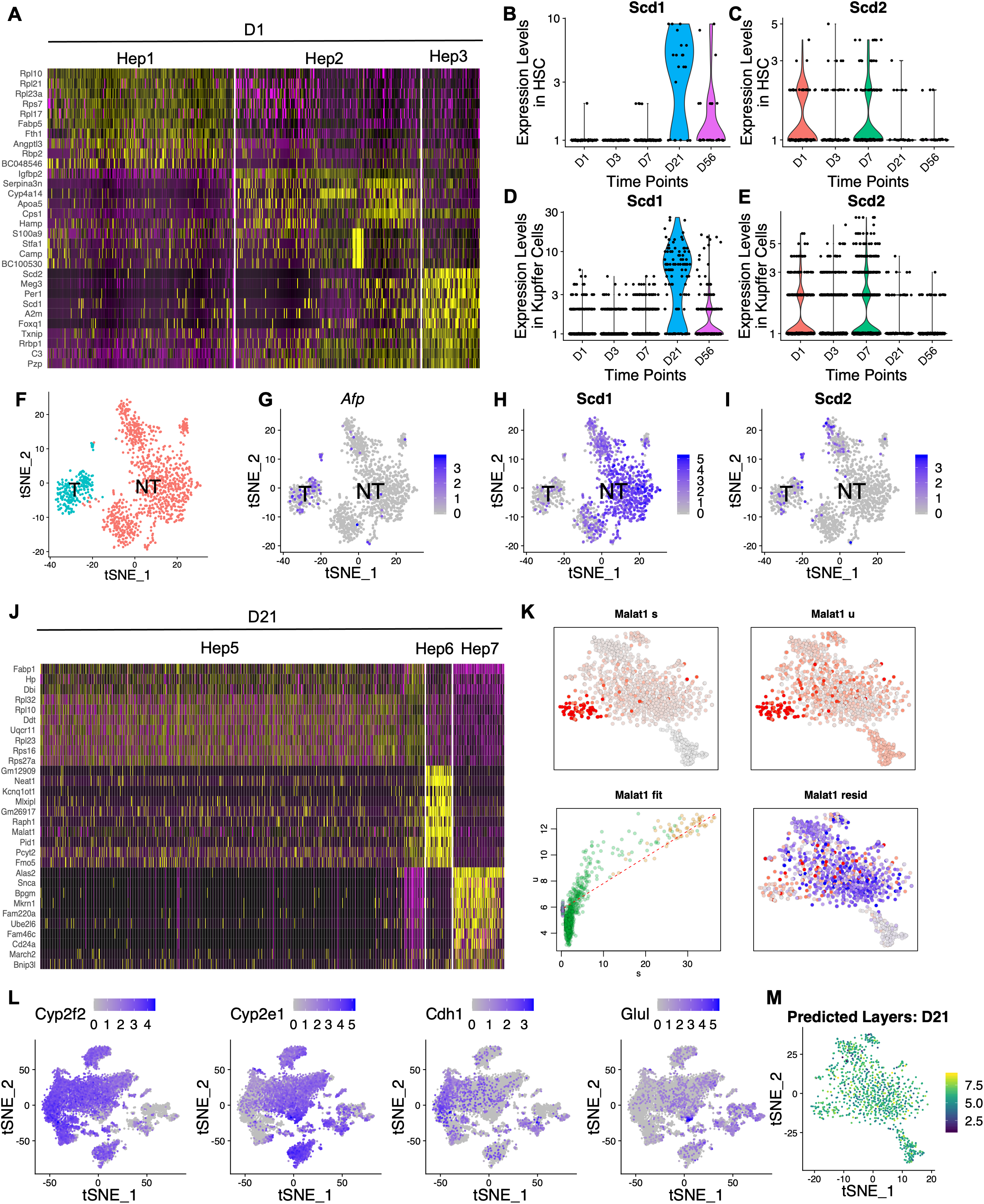
Additional plots associated with hepatocyte subpopulations, related to Figure 2. (A). Heatmap displaying top-ranked marker genes in each cluster identified in Figure 2A (D1). (B-C). Violin plots displaying *Scd1* (B) and *Scd2* (C) expression in HSCs from D1, D3, D7, D21 and D56. (D-E). Violin plots displaying *Scd1* (D) and *Scd2* (E) expression in Kupffer cells from D1, D3, D7, D21 and D56. (F). tSNE map of tumor cells (T) and non-tumor hepatocytes (NT) from Myc-induced mouse HCC model (unpublished data). (G). tSNE map displaying *Afp* expression of cells from Figure S1F. Verified T and NT cell clusters. (H-I). tSNE map displaying *Scd1* (H) and *Scd2* (I) expression in cells from Figure S1F. (J). Heatmap displaying top-ranked marker genes in each cluster identified in Figure 2D (D21). (K). *Malat1* expression pattern obtained with RNA velocity analysis: (s) tSNE map of spliced expression; (u) tSNE map of unspliced expression; (fit) phase portraits showing *Malat1* was induced in Hep6 (colors indicate assigned hepatocyte subpopulations in Figure 2D); (resid) tSNE map of residuals. (L). tSNE map displaying indicated zonation marker expression in hepatocytes from Figure 1D. (M). Predicted layers visualized on tSNE map from Figure 2D (D21).

**Figure S2.**
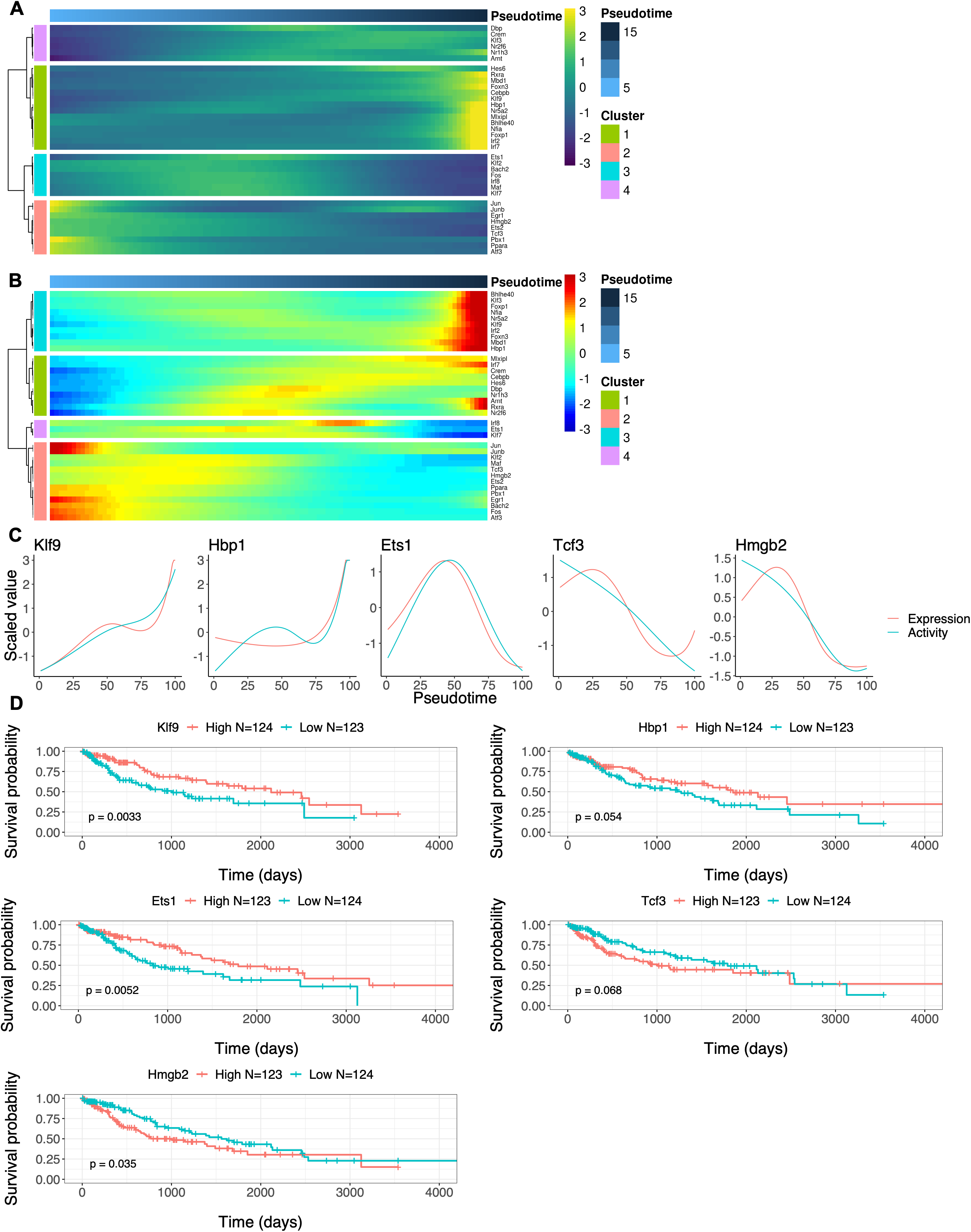
Selected TFs involved in hepatocyte postnatal development and their implication in HCC, related to Figure 3. (A). Heatmap displaying activities of selected TFs along inferred pseudotime in Figure 3B. (B). Heatmap displaying expression of genes encoding selected TFs along inferred pseudotime in Figure 3B. (C). Fitted plot of indicated TF scaled expression and activity values along pseudotime. The inferred pseudotime from Figure 3B was stretched from 0 to 100. (D). Kaplan-Meier survival analysis of HCC patients from TCGA database. Patient samples were divided into High, Mid and Low groups, based on their mRNA expression levels of indicated gene. Mid group was not included in survival analysis. Genes are related to corresponding TFs in Figure S2C.

**Figure S3.**
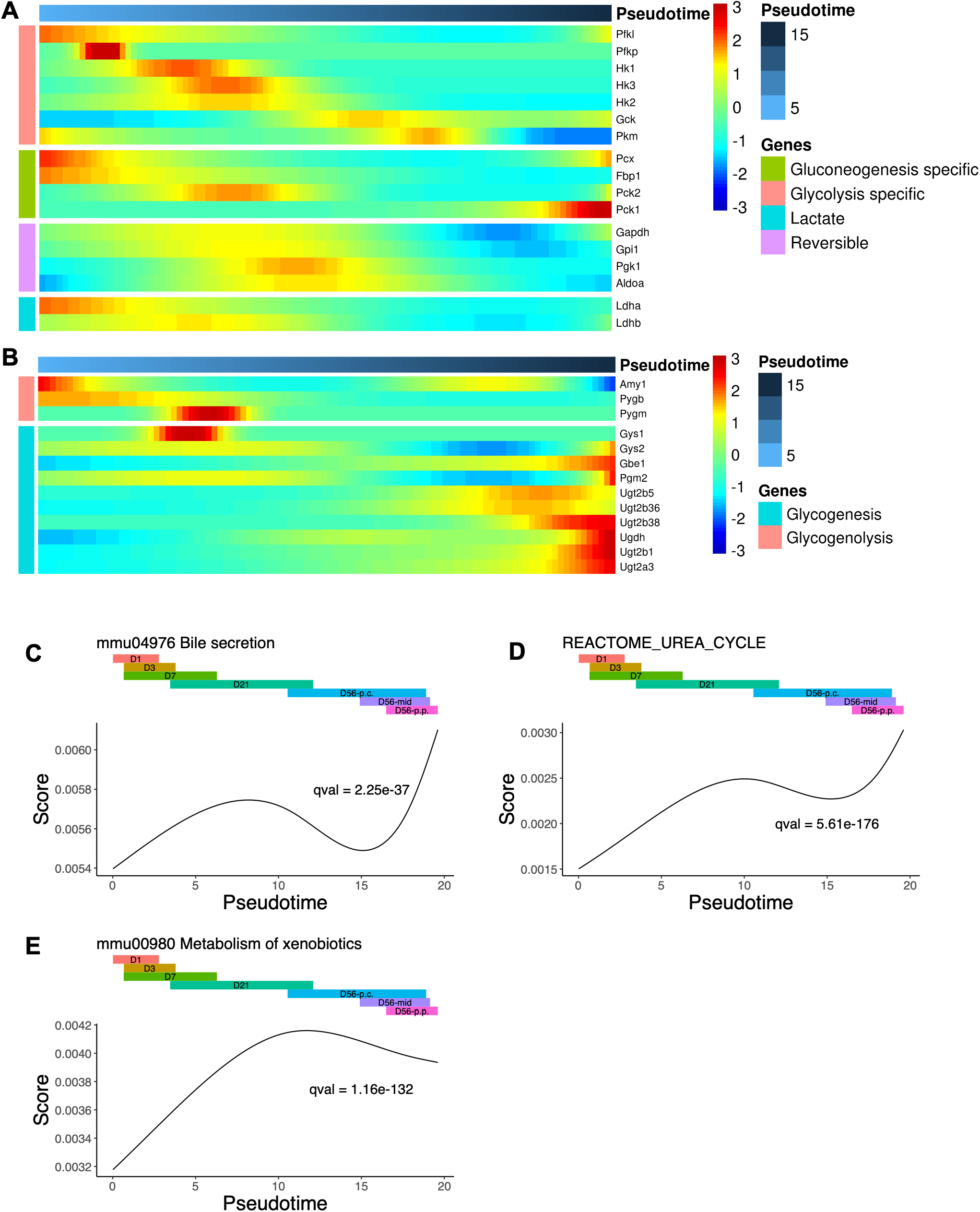
Additional plots associated with metabolic changes in hepatocytes during postnatal development, related to Figure 3. (A). Heatmap displaying genes from pathway “mmu00010 Glycolysis/Gluconeogenesis” with differential expression along inferred pseudotime in Figure 3B. Genes were clustered into four clusters: genes encoding gluconeogenesis-specific enzymes, genes encoding glycolysis-specific enzymes, genes encoding lactate synthesis-related enzymes and genes encoding reversible enzymes. (B). Heatmap displaying genes from pathway “mmu00500 Starch and sucrose metabolism” with differential expression along inferred pseudotime in Figure 3B. Genes were clustered into two clusters: genes encoding glycogenesis-specific enzymes and genes encoding glycogenolysis-specific enzymes. (C-E). Fitted plots of enrichment scores for indicated pathways along inferred pseudotime in Figure 3B.

**Figure S4.**
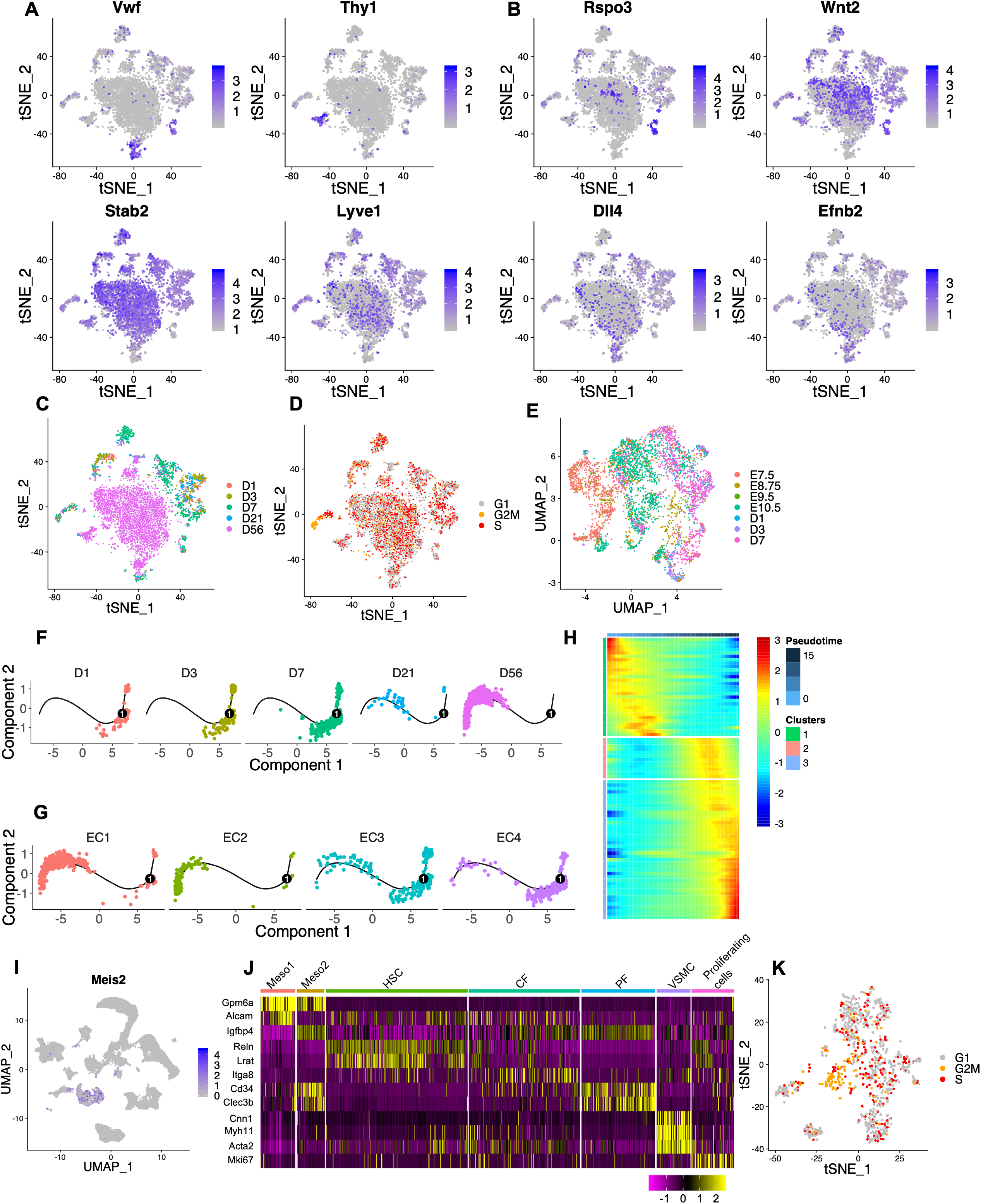
Additional plots associated with endothelial cells from all time points, related to Figure 4. (A). tSNE map displaying indicated endothelial marker expression in cells from Figure 4A. (B). tSNE map displaying indicated endothelial zonation marker expression in cells from Figure 4A. (C). tSNE map of cells from Figure 4A. Colors indicate time points. (D). tSNE map of cells from Figure 4A. Colors indicate predicted cell cycle phases. (E). (E). UMAP visualization of cells from Figure 4C. Colors indicate time points. (F-G). Pseudotime analysis of endothelial cells from D1, D3, D7, D21 and D56 with Monocle 2. Inferred pseudotime was shown in Figure 4F. Colors indicate time points (F) or endothelial cell subpopulations (G). (H). Heatmap representing trends of differentially expressed genes as a function of inferred pseudotime in Figure 4F. (I). UMAP visualization displaying expression level of *Meis2* in all cells included in this analysis (Figure 1A). (J). Heatmap displaying expression of selected markers in mesenchymal cell types identified in Figure 4H. (K). tSNE map of cells from Figure 4H. Colors indicate predicted cell cycle phases.

**Figure S5.**
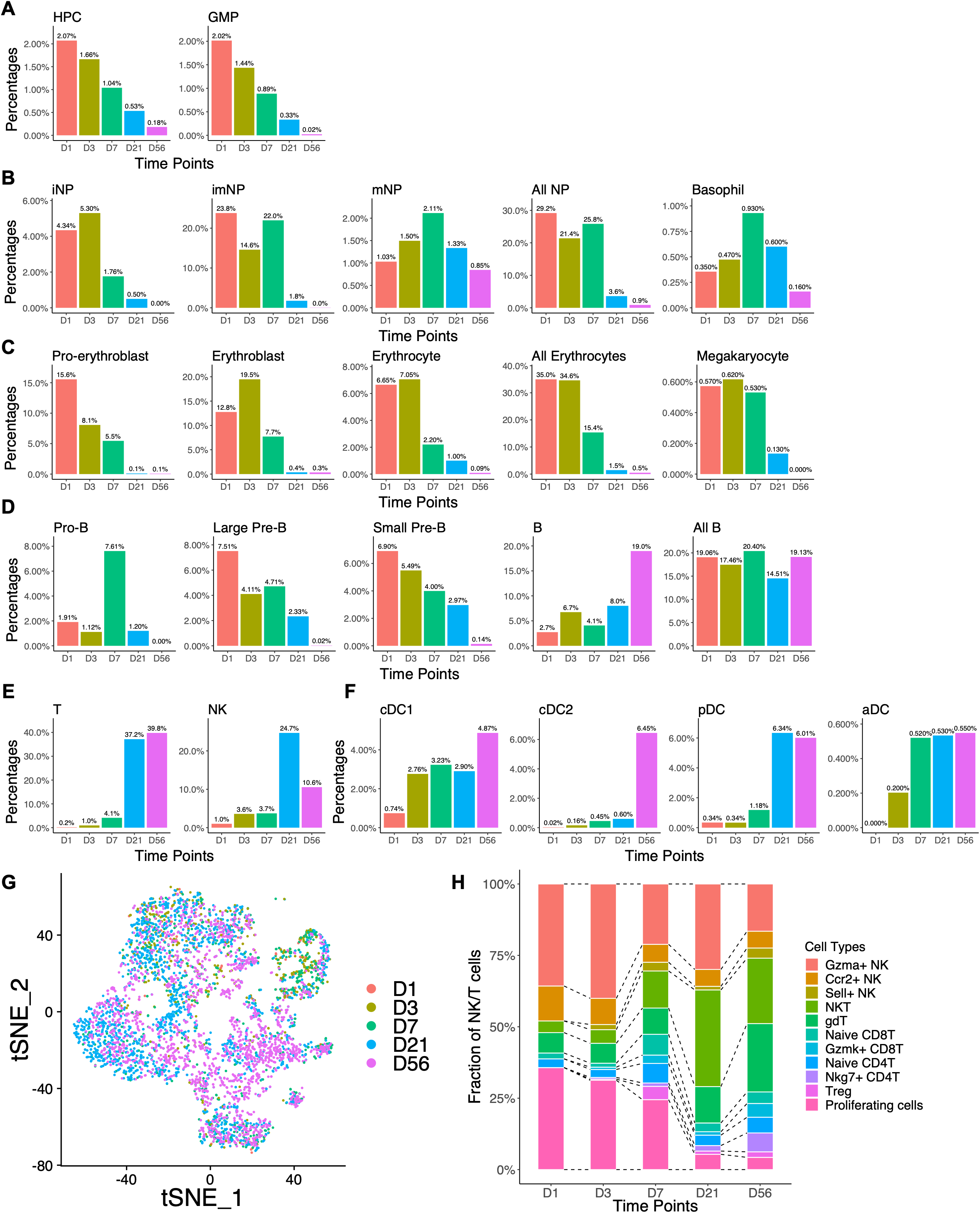
Immune cell subpopulations identified in postnatal liver, related to Figure 5. (A-F). The percentage of indicated cell type out of total immune cell population at each time point, related to Figure 5B. (G). tSNE map of cells from Figure 5C. Colors indicate predicted time points. (H). Time point composition of each T or NK cell type identified in Figure 5C.

**Figure S6.**
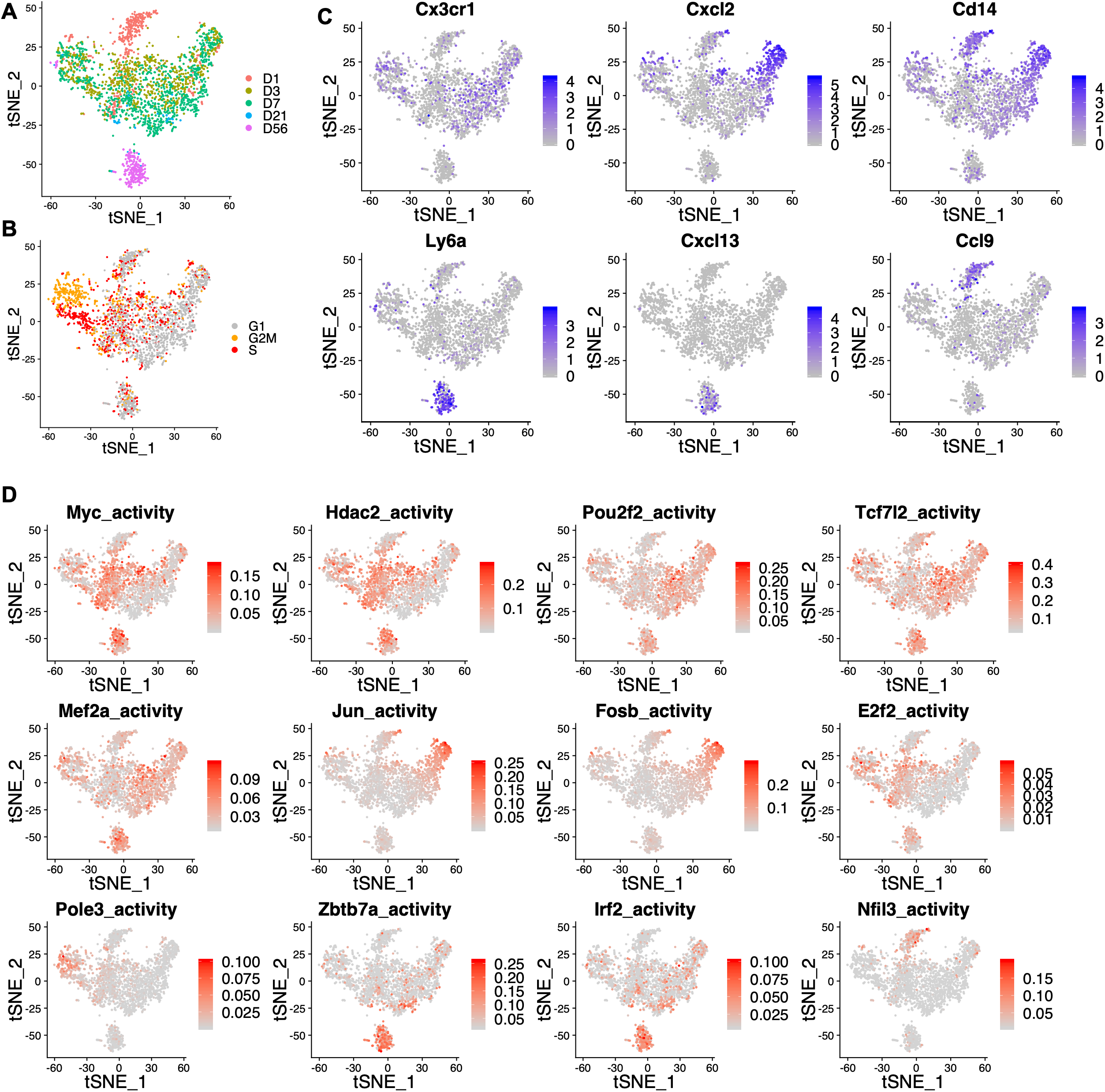
Kupffer cells subpopulation identified in postnatal liver, related to Figure 6. (A). tSNE map of cells in Figure 6C. Colors indicate time points. (B). tSNE map of cells in Figure 6C. Colors indicate predicted cell cycle phases. (C). tSNE map displaying expression levels of selected marker genes for Kupffer cell subpopulations identified in Figure 6C. (D). tSNE map displaying activities of selected TFs for Kupffer cell subpopulations identified in Figure 6C.

**Figure S7.**
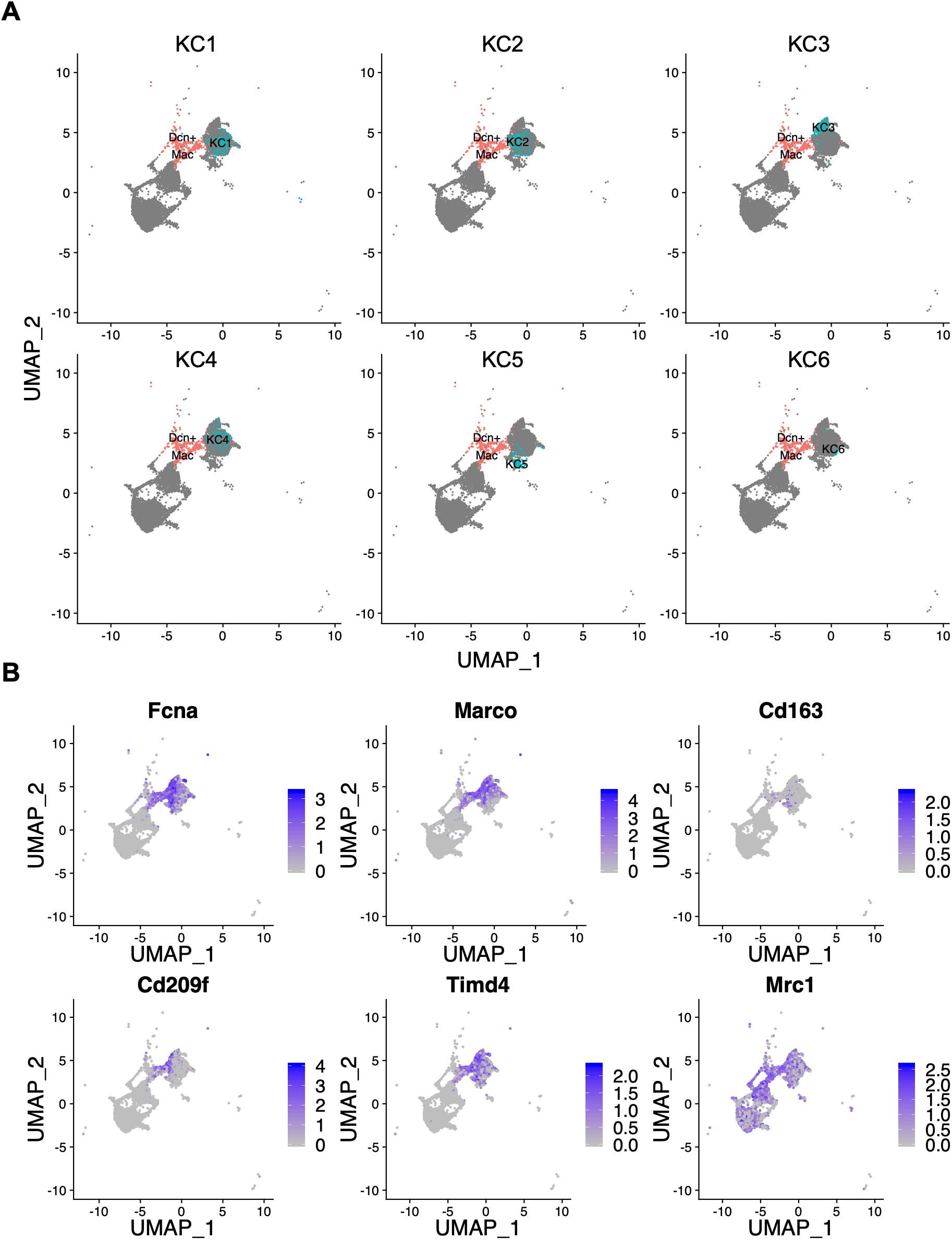
Dcn^+^ Mac cell identified at D7, related to Figure 6. (A). UMAP visualization of cells in Figure 6I. Colors indicates Kupffer cell subpopulation and Dcn^+^ Mac cells. (B). UMAP visualization displaying expression of marker genes of asterisk cluster labelled in Figure 6J.

## STAR METHODS

### KEY RESOURCES TABLES

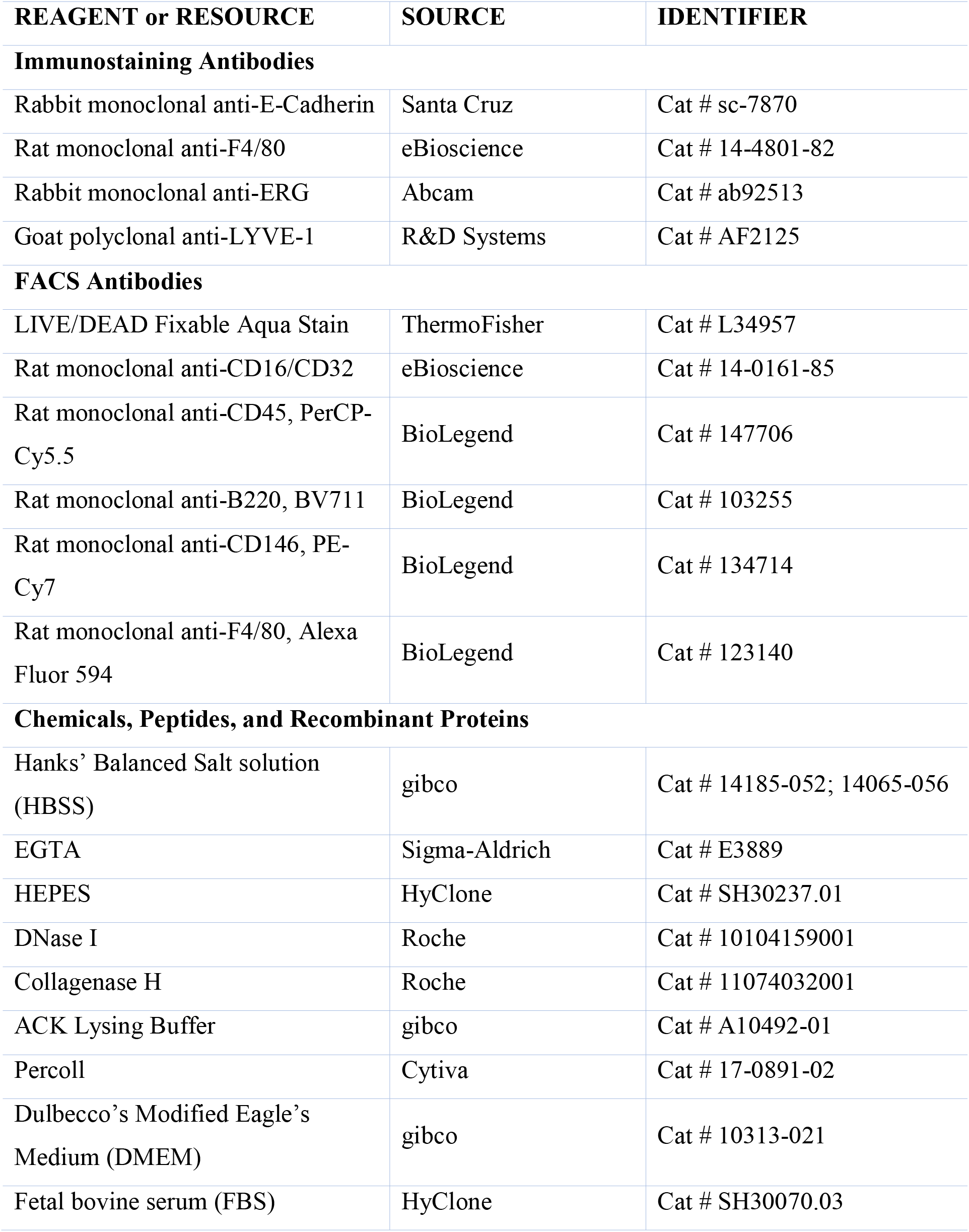
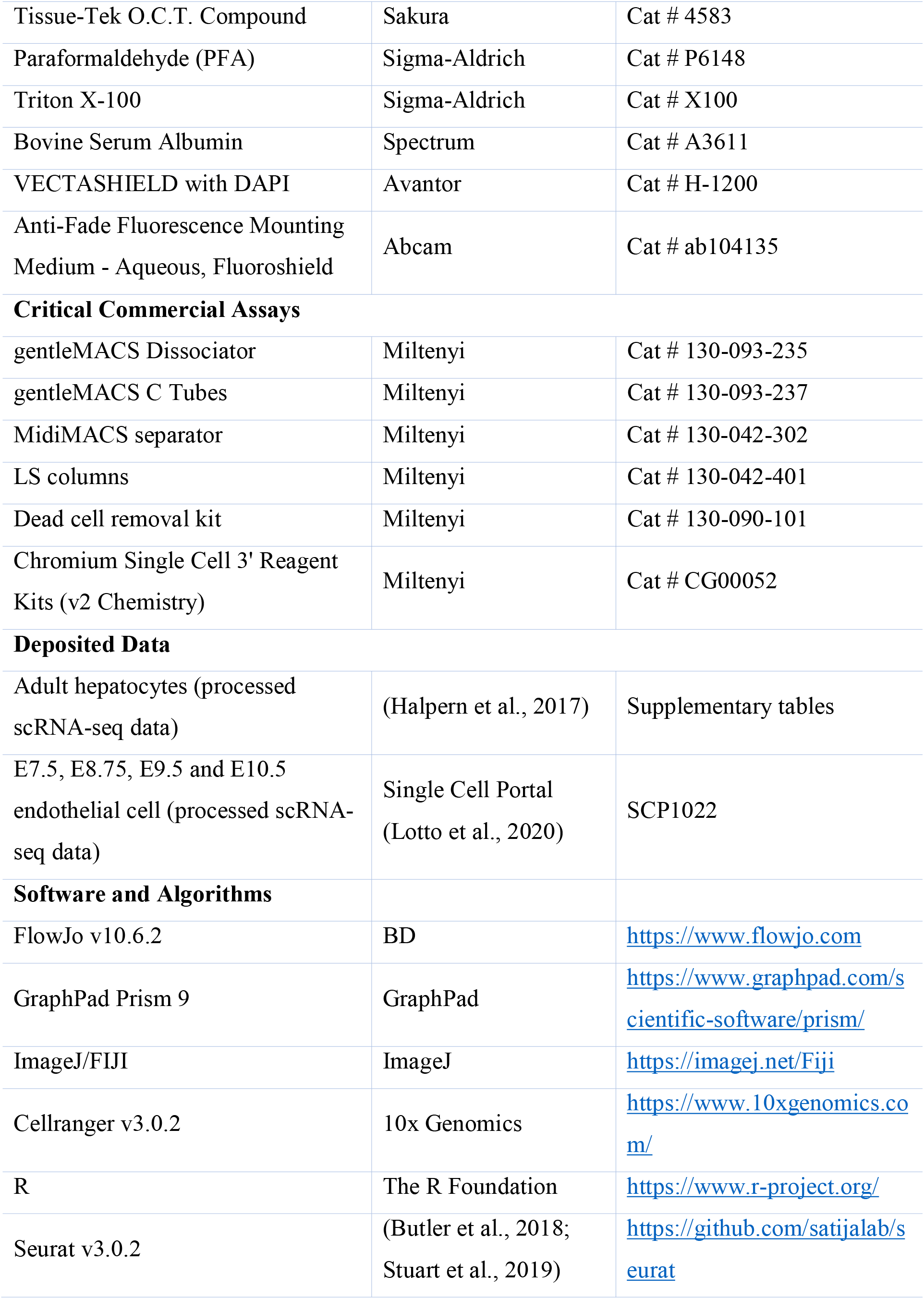
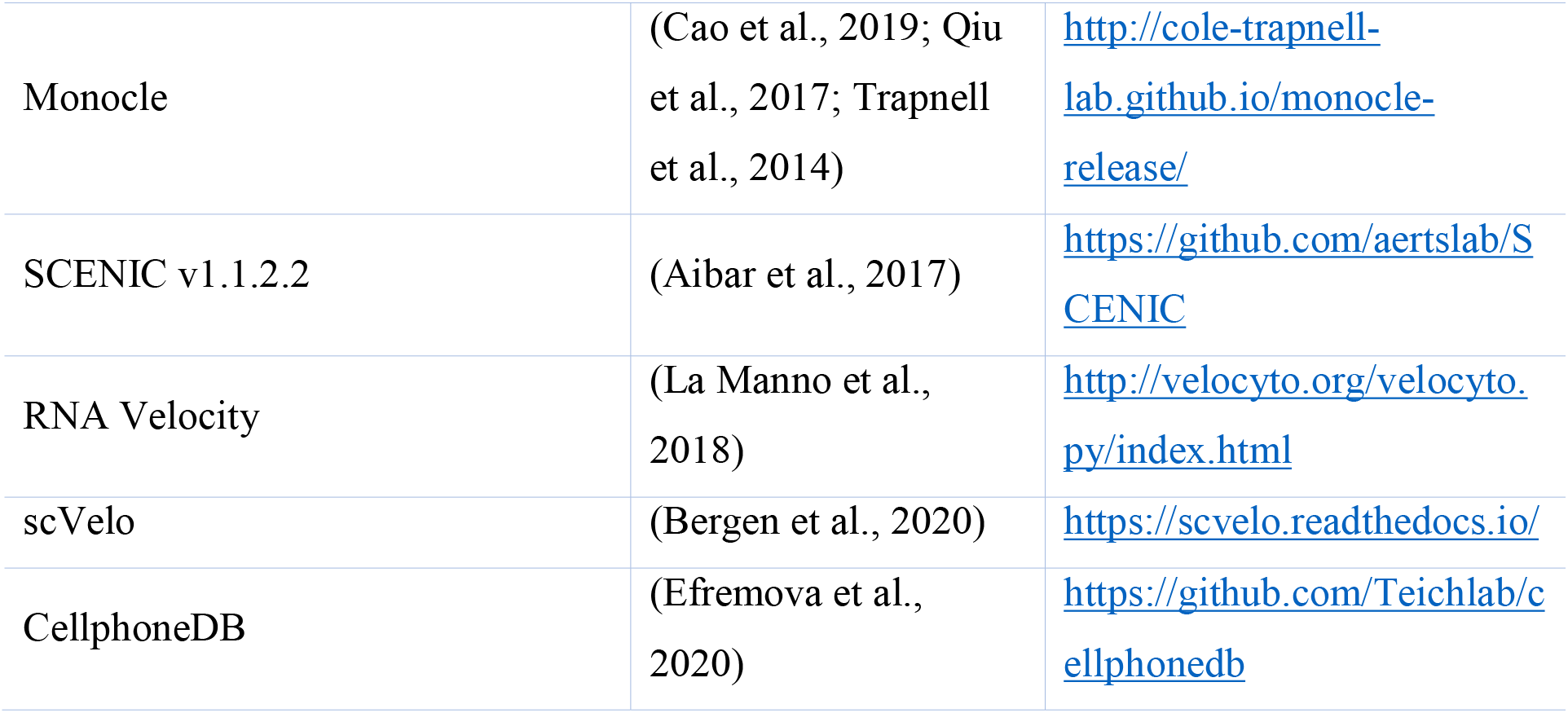

### RESOURCE AVAILABILITY

#### Lead Contact

Further information and requests for resources and reagents should be directed to and will be fulfilled by the Lead Contact, Gen-Sheng Feng.

#### Materials Availability

This study did not generate new unique reagents.

#### Data and Code Availability

The accession number for the raw data and processed data reported in this paper is GSE171993

### EXPERIMENTAL MODEL AND SUBJECT DETAILS

#### Mice

All animals used in this study were in C57BL/6J background. The animal protocol (S09108) was approved by the IACUC at the University of California San Diego.

### METHOD DETAILS

#### Liver cell isolation for scRNA-seq

For neonatal samples (D1, D3 and D7), 4 male mice were included in each time point. Decapitation was used for neonate sacrifice because they are resistance to hypoxia at this age. First, fetal livers were removed from the body, cut into small pieces and rinsed using Ca^2+^-free HBSS to reduce blood. Next, liver pellets were transferred into HBSS buffer with collagenase H and DNase I, further dissociated with gentleMACS program *m_liver_03* and then incubated at 37℃ for 30 minutes. After incubation, the additional gentleMACS program *m_liver_04* was used for more complete dissociation. Liver was passed through a 100 *µ*m cell strainer. Cells were then centrifuged at 50 g for 3 minutes, in order to separate hepatocytes (in pellets) from non-parenchymal cells, NPCs (in suspension). In non-parenchymal cell suspension, ACK buffer was added to lyse red blood cells. Hepatocytes and NPCs were further washed with PBS separately, resuspended in DMEM with 10% FBS, counted with hemocytometer and mixed by original ratios of cell numbers.

For D21 and D56 samples, 2 male mice were included for each time point. Mice were first sacrificed using CO_2_. Liver was perfused using a 2-step method with Ca^2+^-free HBSS buffer and then with collagenase H in HBSS buffer containing Ca^2+^, gently minced and passed through a 100 *µ*m cell strainer. Cells were centrifuged at 50 g for 3 min, in order to separate hepatocytes (in pallets) from NPCs (in suspension). To remove dead cells and debris, the pelleted hepatocytes were resuspended in 45% Percoll and centrifuged at 50 g for 10 min, without brake. Meanwhile, the ACK buffer and Dead Cell Removal Kit were used to lyse red blood cells and eliminate dead cells in non-parenchymal cells. Hepatocytes and NPCs were further washed with PBS separately, resuspended in DMEM with 10% FBS and counted with hemocytometer.

Liver single cell preparations were made within the same four-hour period of a day for collection of samples at all five time points.

#### Single cell library construction and sequencing

The isolated single cells were immediately loaded onto 10x Chromium Controller, and then partitioned into nanoliter-scale Gel Beads-In-Emulsion (GEMs). Cells in GEMs were lysed, and RNAs released from cells were immediately captured by barcoded beads in the same GEMs, followed by reverse transcription, amplification, fragmentation, adaptor ligation and index PCR. Libraries were constructed using Chromium Single Cell 3’ Reagent Kits (V2 chemistry, 10x Genomics). Sequencing was performed on Illumina HiSeq 4000 at IGM Genomics Center, UCSD, with the following read length: Read 1, 26 bp, including 16 bp cell barcode and 12 bp unique molecular identifier (UMI); Read 2, 98 bp transcript insert; i7 sample index, 8 bp.

#### scRNA-seq data pre-processing, dimensionality reduction and clustering

Sequenced reads were aligned to mouse reference genome GRCm38 using CellRanger package (v3.0.2). All libraries were then aggregated for batch effect correction and sequencing depth normalization. An expression matrix including all cells from D1 to D56 livers was generated, with each row representing a gene and each column representing a cell, and then loaded into the R package, Seurat. Next, low quality cells and genes were filtered for downstream analysis. In brief, genes expressed in less than 3 cells were removed; cells failed to meet following criteria were removed: 1) the number of genes detected in each cell should be more than 200 but less than 6500; 2) the UMIs of mitochondrial genes should be less than 10% of total UMI. A total of 52834 cells and 24057 genes passed the filter. After filtering, the raw expression matrix was normalized by the total expression, multiplied by scale factor 10,000, and log transformed. Next, we regressed on total numbers of UMIs per cell as well as mitochondrial gene percentages. The z-scored residuals calculated by normalization and scaling were stored for downstream analysis. We then performed dimensionality reduction and unsupervised clustering using Seurat functions. In brief, PCA was performed first, and 75 PCs were loaded for UMAP and tSNE dimensionality reduction. To find clusters, the same PCs were imported into FindClusters, a SNN graph-based clustering algorithm, with resolution = 3.0. Next, the Wilcoxon Rank Sum test was performed to identify markers (FDR < 0.05) in each cluster. Cell types were then assigned based on these markers manually.

Further, to find subpopulations in hepatocyte, endothelial cells, mesenchymal cells, NK cells, T cells and Kupffer cells, cells from corresponding groups were segregated and re-analyzed for higher resolution.

For combined analysis of embryonic (Lotto et al., 2020) and fetal (this work) endothelial cells, an additional dataset integration using Seurat was perform to remove batch effect.

#### Zonation profile prediction of hepatocytes

We predicted spatial layers (zonation) of hepatocytes using linear regression method. The training dataset was downloaded from previously published work (Halpern et al., 2017), with processed expression matrix and zonation information for each cell. We normalized, scaled and integrated this dataset with our hepatocyte subpopulations to remove batch effect. Next, the Wilcoxon Rank Sum test was performed to identify markers (FDR < 0.05) in each layer. We combined markers for 9 layers to make a zonation signature with 210 genes in total. With zonation signature gene set, the linear model was trained using original dataset (Halpern et al., 2017) and then used to predict layer for single hepatocyte in this study. The predicted layer was then normalized from 1 to 9, representing spatial distribution of hepatocytes from pericentral to periportal area.

#### Trajectory analysis

We performed trajectory analysis using different methods listed below, depending on different situations and purposes.

##### RNA velocity analysis

As described in original publication (La Manno et al., 2018), this algorithm was designed based on a simple model for transcriptional dynamics, in which the RNA velocity was estimated from spliced and un-spliced mRNA, and then used to predict the future state of individual cells. We first generated loom files of spliced and un-spliced reads counts from 10x samples using velocyto *run10x* pipeline. We then visualized RNA velocity on specified tSNE or UMAP embedding. For gene of interest, we also plotted (1) spliced or (2) un-spliced counts on the same embedding, (3) phase portraits against steady-state (un-spliced levels on y-axis above steady-state represented gene of interest being induced, while un-spliced levels below steady-state represented being repressed), and (4) residual levels *u*, with positive (red) and negative (blue) values indicating induced and repressed, respectively, embedded onto the same dimensionality reduction.

##### Monocle

This method was used to perform trajectory analysis and calculate pseudotime for a relatively small group of cells. In brief, we selected differentially expressed genes between different time points as ordering genes and applied DDRTree, which learned the structure of dataset manifold as developmental trajectory and ordered cells onto that manifold with a calculated pseudotime starting from a given root cell. Next, we selected genes significantly changed as a function of pseudotime along trajectory. In brief, for each gene, the expression level was fitted with natural spline (sm.ns) to describe the smooth change of this gene along trajectory. The smooth values were then employed to fit a negative binomial model via vglm() function from VGAM package. This was the full model. Next, to test for significance, a chi-square ratio test was performed to compare this full model against null model. The whole process was wrapped in *differentialGeneTest* function in monocle. We adapted this method here and also for analysis described below. Finally, we made heatmap with significantly changed genes, clustered them by pseudotemporal expression pattern and then performed pathway enrichment analysis with gene sets from Msigdb v7.0 (Subramanian et al., 2005).

##### Monocle 3

For a much larger dataset, we performed trajectory analysis for integrated embryonic and fetal endothelial cells using monocle 3 with UMAP visualization for a better performance.

##### Force-directed graph analysis

This method was performed to visualize connections between immature hematopoietic cells and their progenitor cells, with pseudotime calculated (not shown). The code was adapted from published work on fetal liver hematopoiesis (Popescu et al., 2019).

#### Gene regulatory network analysis

SCENIC (Single Cell Regulatory Network Inference and Clustering) was performed to identify upstream regulators important for hepatocyte, LSEC and Kupffer cell development individually. With segregated raw expression matrix for cell type of interest, we further filtered for genes expressed at least with a count of 3 in 1% of all cells and genes found in RcisTarget’s mouse databases (mm9-500bp-upstream-7species.mc9nr.feather, mm9-tss-centered-10kb-7species.mc9nr.feather). Next, the filtered expression matrix was normalized as log2(exprMat + 1). SCENIC was then performed to identify and score regulons (TFs) based on the expression of their regulated target genes. With regulon scores, we adopted differential expression function from Monocle to test for TFs with significantly changed activities as a function of pseudotime. We fitted regulon score with gamma distribution instead and also performed chi-square ratio test for significance. A similar heatmap was plotted for TF activities using regulon scores. Also, for a TF of interest, we fitted natural spline to its scaled expression values (Expression) and regulon scores (Activity) and plotted the smooth change of said TF along pseudotime, which was stretched from 0 to 100.

#### Single cell pathway enrichment

We collected metabolism-related pathways from Kyoto Encyclopedia of Genes and Genomes (KEGG) and Msigdb. To find metabolic pathways significantly changed during postnatal development, we first calculated enrichment score for individual cell i of each gene set j. The score was defined as follows:

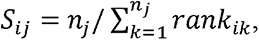

where n_j_ represented the length of gene set j; rank_ik_ represented the rank of cell i ordered by expression levels of gene k (in gene set j) over all cells included in this analysis. Next, for each gene set j, we ordered cells along trajectory and tested if S_ij_ changed significantly as a function of pseudotime. The statistical test method was also adapted from monocle as described above. We fitted gene set enrichment scores with gamma distribution and performed chi-square ratio test for significance. Similarly, we plotted smooth scores after fitting the natural spline.

#### Cell-cell interaction analysis

We performed CellPhoneDB with our dataset to identify important ligand-receptor interactions. All cell types were included for the analysis at each time point. The genes encoding ligands and receptors were kept only if they were expressed by at least 10% of cells. The significance of an interaction was calculated by permutation test with iterations = 1000.

#### TCGA survival analysis

The TCGA dataset of HCC was downloaded from TCGA-LIHC project, including processed results of RNA-sequencing and clinical information for 371 patients. For each gene of interest, all samples were divided into three groups based on gene expression levels: Low (patients with expression levels lower than 33.3% percentile); High (patients with expression levels higher than 66.7% percentile); Mid (other patients). Survival analysis was performed using survival and survminer R package, including the Low and High groups divided based on gene of interest.

#### Immunostaining

To prepare fresh frozen tissue sections, all liver tissues were collected at indicated time points and were immediately embedded Tissue-Tek O.C.T compound (Sakura) and stored at -80℃ . Fresh frozen tissue sections were fixed by cold acetone overnight, followed by cold 4% PFA overnight. After primary and secondary antibody incubation, fresh frozen tissue sections were mounted with VECTASHIELD mounting medium with DAPI or Anti-Fade Fluorescence Mounting Medium (for confocal microscope). For single-cell resolution, we checked slides under Leica SP8 Confocal with Lightning Deconvolution at microscopy core, University of California, San Diego.

#### Flow cytometry analysis

Non-parenchymal cells were isolated from liver as described above (see **Liver cell isolation for scRNA-seq**). Cells were first stained for LIVE/DEAD fixable Aqua, anti-CD16/CD32 for Fc blocking, and then panel antibodies. Cells were fixed using 1% PFA in PBS overnight at 4℃, resuspended in PBS, and analyzed on BD LSRFortessa X-20 Cell Analyzer at UCSD Human Embryonic Stem Cell Core facility at Sanford Consortium for Regenerative Medicine. Data were then analyzed using FlowJo 10.6.2.

#### Statistical analysis

The statistical analysis was performed using R or GraphPad Prism 9 (for FACS analysis only). The details of test and significance are specified in figure legends.

## Notes

### Competing Interest Statement

The authors have declared no competing interest.

